# A novel in vitro tubular model to recapitulate features of distal airways: The bronchioid

**DOI:** 10.1101/2023.12.06.569771

**Authors:** Elise Maurat, Katharina Raasch, Alexander M. Leipold, Pauline Henrot, Maeva Zysman, Renaud Prevel, Thomas Trian, Tobias Krammer, Vanessa Bergeron, Matthieu Thumerel, Pierre Nassoy, Patrick Berger, Antoine-Emmanuel Saliba, Laetitia Andrique, Gaëlle Recher, Isabelle Dupin

## Abstract

**Background:** Airflow limitation is the hallmark of obstructive pulmonary diseases, with the distal airways representing a major site of obstruction. Although numerous *in vitro* models of bronchi already exist, there is currently no culture system for obstructive diseases that reproduces the architecture and function of small airways. Here, we aimed to engineer a model of distal airways to overcome the limitations of current culture systems.

**Methods:** We developed a so-called bronchioid model by encapsulating human bronchial adult stem cells derived from clinical samples in a tubular scaffold made of alginate gel.

**Results:** This template drives the spontaneous self-organization of epithelial cells into a tubular structure. Fine control of the level of contraction is required to establish a model of the bronchiole, which has a physiologically relevant shape and size. 3D imaging, gene expression and single-cell RNA-seq analysis of bronchioids made of bronchial epithelial cells revealed tubular organization, epithelial junction formation and differentiation into ciliated and goblet cells. Ciliary beating is observed, at a decreased frequency in bronchioids made of cells from COPD patients. The bronchioid can be infected by rhinovirus. An air-liquid interface is introduced that modulates gene expression.

**Conclusion:** Here, we provide a proof of concept of a perfusable bronchioid with proper mucociliary and contractile functions. The key advantages of our approach, such as the air-liquid interface, lumen accessibility, recapitulation of pathological features and possible assessment of clinically relevant endpoints, will make our pulmonary organoid-like model a powerful tool for preclinical studies.

## Introduction

Chronic obstructive diseases such as chronic obstructive pulmonary disease (COPD) and asthma are major public health challenges characterized by airflow limitation. There is a lack of specific curative treatments for these diseases, at least because of the paucity of integrative approaches for deciphering their pathophysiology, especially for the distal airways. On the one hand, animal and especially rodent models are widely used for preclinical studies, but their airways differ anatomically, histologically and physiologically from those of humans, and they do not fully recapitulate disease hallmarks [1]. To date, they poorly predicted which drugs will help treat patients with chronic obstructive diseases. On the other hand, the lung has a complex structure leading to a peculiar stiffness and 3D conformation that is difficult to model in vitro, highlighting the need to create adequate complex cellular models.

Various strategies have been proposed to model human bronchi *in vitro* [2]. 2D culture of bronchial epithelial cells at the air-liquid interface (ALI) allows the reconstitution of a fully differentiated epithelium with functional cilia, with easy access to the apical and basal sides [3, 4], but it does not reproduce the three-dimensional architecture essential for properly modelling the airways. 3D bioprinting is a promising method for building tissue with a relevant geometry [5, 6], especially with the recent development of suitable bioinks, such as those composed of alginate and decellularized tissue, enabling the printing of a tubular structure [7]. However, the length of the tubes obtained is limited, and 3D printing of the last branches of the bronchial tree remains below the resolution limit of current bioprinting techniques. The elegant microfluidic approach has been used to design so-called “airway-on-a-chip” [8, 9], providing unique advantages such as the ability to perfuse air, particles or nebulize drugs, to incorporate immune cells and the potential application of cyclic stretching to reproduce breathing movements. While the microengineered scaffold restricts morphogenesis to a predefined geometry, it does not completely match the exact topology of a bronchi and does not offer a permissive and remodelable environment.

Other studies, starting from Benali *et al*.[10] to more recent ones [11–13] have favoured the 3rd dimension with “bronchospheres”, in which bronchial epithelial cells are embedded in a gel and self-organize into spheres with a liquid-filled lumen [14]. These self-renewing organoids exhibit ciliary activity and mucus production. In most of the bronchospheres, the apical side of the cells faces the inside of the organoid, limiting access to the lumen, although protocols to reverse airway organoid polarity have been developed [15, 16]. This closed and spherical architecture also impairs the removal of apical side-released dead cells inside and the introduction of an air-liquid interface. Finally, the formation of invasive tubular structures from lung organoids has been observed [17], but this process remains stochastic and rather limited.

The Cellular Capsule Technology, adapted from a co-extrusion microfluidic technique, has been used to encapsulate cells and extracellular matrix proteins in a tubular scaffold made of alginate gel [18]. Cell-laden microfibres have been generated with various cell types, such as endothelial cells, epithelial cells, fibroblasts, skeletal muscle cells and nerve cells [18]. When co-encapsulated, endothelial and vascular smooth muscle cells self-organize into tubular structures that model vessels [19]. Although epithelial cell lines such as the human hepatoma G2 (HepG2) cell line, the mouse pancreatic beta cell (MIN6m9) cell line, the cervical cancer Henrietta Lacks (HeLa) cell line, the Madin-Darby Canine Kidney (MDCK) cell line and the EpH4-J3B1A mammary gland epithelial cell line have been previously successfully used [18, 20], the behaviour of primary bronchial epithelial cells in this culture system has never been investigated.

Here, we aimed to construct a tubular organoid-like model of distal airways with human primary bronchial epithelial cells. By using a tubular template made of alginate using the Cellular Capsule Technology, we were able to drive the spontaneous self-organization of lung epithelial cells to obtain a 3D “bronchioid” model with a physiologically relevant shape and size. Importantly, the tubular geometry allows lumen access to create an air-liquid interface.

## Methods

### Patients

Macroscopically normal lung resection material for the purification of basal epithelial cells was obtained from a cohort of patients requiring thoracic lobectomy surgery for nodules or cancer (pN0) (*i.e.*, TUBE, sponsored by the University Hospital of Bordeaux, which includes its own local ethics committee (CHUBX 2020/54)). According to French law and MR004 regulations, patients received an information form, allowing them to refuse the use of their surgical samples for research. The research was conducted according to the principles of the World Medical Association Declaration of Helsinki. A total of 11 COPD patients with a clinical diagnosis of COPD according to the GOLD guidelines [21] and 18 non-COPD patients with normal lung function (*i.e.*, FEV1/FVC > 0.70) were enrolled from the University Hospital of Bordeaux (Table 1). A total of 5 other patients were also enrolled from the TUBE cohort for the comparison of bronchioids to classical ALI culture (Table E1). All clinical data were collected at the Clinical Investigation Center (CIC1401) of the University Hospital of Bordeaux.

**Table 1:** Patient characteristics for bronchioid generation.

|  | **Non-COPD** | **COPD** | **P value** |
| --- | --- | --- | --- |
|  | **patients** | **patients** |  |
| n | 18 | 11 |  |
| Age (years) | 67.3 ± 8.6 | 67.0 ± 5.2 | 0.92 |
| Sex (Male/Female) | 8/10 | 5/6 | >0.99 |
| Current/Former/Non smokers | 5/10/3 | 2/8/1 | 0.64 |
| Pack years (no.) | 26.7 ± 17.8 | 42.7 ± 33.6 | 0.22 |
| **PFT** |  |  |  |
| FEV1 (% pred.) | 90.9 ± 29.6* | 73.2 ± 22.8* | 0.08 |
| FEV1/FVC ratio (%) | 75.7 ± 10.5* | 61.8 ± 13.8** | <0.0001 |

### Bronchial epithelial cell culture

Human basal bronchial epithelial cells (BECs) were derived from bronchial specimens as described previously [22]. Bronchial epithelial tissue was cultured in PneumaCult™-Ex medium (Stemcell Technologies, Vancouver, Canada) under a water-saturated 5% CO2 atmosphere at 37 °C until basal BECs reached 80-90% confluence. For ALI culture, basal BEC were plated (1.105 cells per well) on tissue culture-treated nucleopore membranes (Ref 3470, 6.5 mm diameter, 0.4 µm pore size, Transwell, Costar) in PneumaCult™-ALI Medium (Stemcell Technologies) applied at the basal side only to establish the ALI. The cells were maintained in culture for 9 or 21 days.

### Microfluidic co-extrusion device and bronchioid formation

The design of a co-extrusion microfluidic device for bronchioid formation has been previously described for vessels [19]. Using this device with a 450 µm-diameter nozzle, we produced hollow alginate tubes filled with basal BECs. Cells were harvested by using 0.05% trypsin-ethylene diamine tetra-acetic acid (EDTA, Fisher Scientific) and resuspended in Matrigel (Corning, New York, USA) and Dulbecco’s modified Eagle’s medium (Sigma-Aldrich, Saint Quentin-Fallavier, France), both of which ranged from 33.5 to 35% (v/v). The cell loading of the core cell suspension (CCS) varied from 30 to 33% (v/v). Cells were then injected into the central zone isolated from a 2% (w/v) alginate solution (AS) (I3G80, Alliance Gums & Industries, Ile de France, France) by a 300 mM intermediate D-sorbitol solution (IS) (Sigma-Aldrich). The experiments were performed using the following injection flow rates: 2 mL/hour (AS), 1 mL/hour (IS), and 1 mL/hour (CCS). With these flow rates, the thickness of the alginate walls was approximately 50 µm, resulting in a 350 µm diameter cell tube (Figure 1A). A 100 mM calcium bath at 37 °C was used to trigger rapid gelation of the alginate gel while avoiding osmotic shock. Immediately after formation (within less than 10 min), the cell-laden tubes were transferred to Pneumacult™ Airway Organoid Seeding Medium (Stemcell Technologies) at 37 °C. After 24 hours, the bronchioids were transferred to Pneumacult™ Airway Organoid Differentiation Medium (Stemcell Technologies) to initiate differentiation. To inhibit Rho-associated protein kinase (ROCK), the Airway Organoid Medium was supplemented with Y-27632 (Sigma-Aldrich) at a concentration of 10 µM. The media were changed daily.

**Figure 1.**
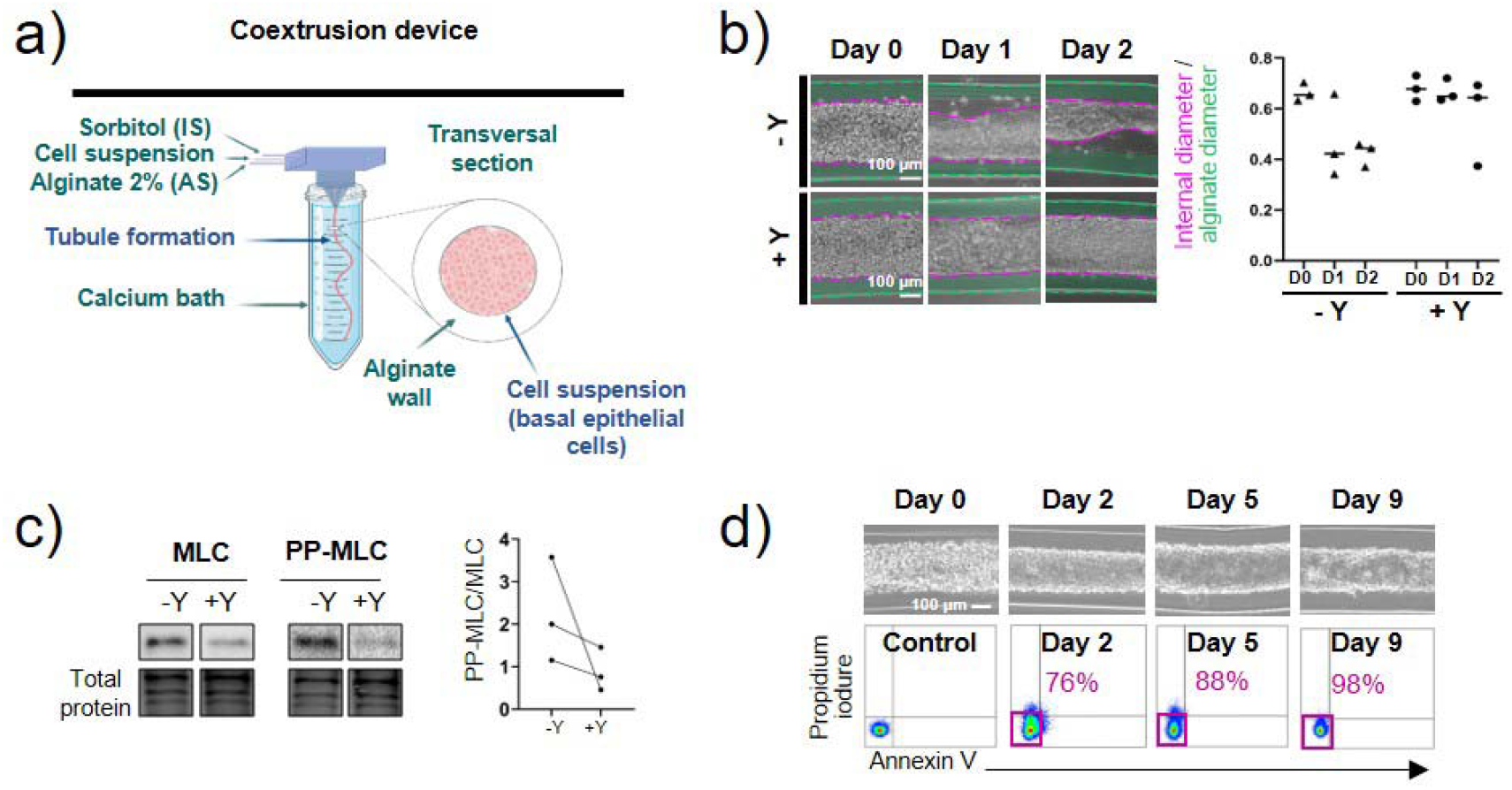
The formation of an alginate tube filled with primary basal epithelial cells is called “bronchioid”. a) Schematic of the device used to generate bronchioids. Created with BioRender.com. The three solutions were injected simultaneously by a computer-controlled pump inside a 3D-printed device soaked in a 100 mM calcium bath. b) Left panel: brightfield images of bronchioids over time, with (+) or without (-) 10 µM Y-27632 (“Y”). The green and pink dotted lines indicate the external (alginate tube) and internal (epithelial tube) limits, respectively. Right panel: quantification of external and internal diameters (at least 10 measurements along the same tube for each condition, n=3 experiments). The medians are represented as horizontal lines. c) Immunoblots and analysis of myosin light chain (MLC) II and double-phospho-MLC (Thr18, Ser19; PP-MLC) expression on day 2 in bronchioid tissue from 3 different donors (n=3). Total protein was used as a loading control. d) Upper panels: brightfield images of bronchioids cultured with 10 µM Y-27632. Lower panels: dot plots showing propidium iodide (PI) fluorescence (y-axis) versus annexin V-allophycocyanin (APC) fluorescence (x-axis) in cells dissociated from bronchioids and analysed by flow cytometry at the indicated time points. The percentages of PI-Annexin V-cells are shown in pink.

### Western blot

The bronchioids were assayed for myosin light chain (MLC) II and double-phospho-MLC (Thr18, Ser19; PP-MLC) levels by Western blotting. Total protein extracts were obtained by transferring the bronchioids to Dulbecco′s phosphate-buffered saline (DPBS, Sigma-Aldrich) for 30 minutes at 37 °C with gentle agitation to dissolve the alginate. After centrifugation, the pellet was suspended in RIPA lysis buffer (Santa Cruz Biotechnology, Dallas, Texas, USA) supplemented with protease inhibitor cocktail and calyculin A (Sigma-Aldrich). These extracts were loaded onto a 4-20% SDS PAGE gel (Bio-Rad, Hercules, California, USA), transferred onto a nitrocellulose membrane and identified with a rabbit anti-target antibody (see Table E2). HRP-coupled anti-rabbit secondary antibody was used for visualization using a ChemiDoc imaging instrument (Bio-Rad). Protein expression was normalized to total protein loading (Stain-free system, Bio-Rad).

### Quantitative real-time PCR

Total RNA was extracted from bronchioids using an RNeasy Kit according to the manufacturer’s instructions (Qiagen, Hilden, Germany). Five hundred nanograms of RNA were reverse-transcribed using the iScript™ Reverse Transcription Supermix (Bio-Rad). cDNA samples were then analysed by qPCR using Perfecta® SYBR® Green SuperMix (Quantabio) through the CFX Connect real-time PCR detection system (Bio-Rad). Primers (*PPIA*, Forward 5′-CGGGTCCTGGCATCTTGT-3′ and Reverse 5′-CAGTCTTGGCAGTGCAGATGA-3′; *RPL13*, Forward 5′-GGGAGCAAGGAAAGGGTCTTA-3′ and Reverse 5′-CACCTGGACAATTCTCCGAGT-3′; *GusB*, Forward 5′-CCATCTGGGTCTGGATCAAAA-3′ and Reverse 5′-TGAAATCGGCAAAATTCCAAAT-3′; *KRT5*, Forward 5′-GCATCACCGTTCCTGGGTAA-3′ and Reverse 5′-GACACACTTGACTGGCGAGA-3′; *TP63*, Forward 5′-CCTCAGGGAGCTGTTATCCG-3′, Reverse 5′-ATACTGGGCATGGCTGTTCC-3′; *SCGB1A1*, Forward 5′-TCCTCCACCATGAAACTCGC-3′ and Reverse 5′-AGGAGGGTTTCGATGACACG-3′; *DNAH5*, Forward 5 - AGAGGCCATTCGCAAACGTA-3′ and Reverse 5′-CCCGGAAAATGGGCAAACTG-3′; *AGR2*, Forward 5′-AAGGCAGGTGGGTGAGGAAATC-3′ and Reverse 5′-GTCGAGAGTCCTTTGTGTCCTT-3′; *MUC5AC*, Forward 5′-TACTCCACAGACTGCACCAACTG-3′ and Reverse 5′-CGTGTATTGCTTCCCGTCAA-3′; *EPCAM*, Forward 5′-CCATGTGCTGGTGTGTGAAC-3′ and Reverse 5′-AACGCGTTGTGATCTCCTTCT-3′; *CDH1* Forward 5′-TTACTGCCCCCAGAGGATGA-3′ and Reverse 5′-TGCAACGTCGTTACGAGTCA-3′;) were purchased from Sigma-Aldrich. mRNA expression was determined using the comparative 2-ΔΔCt method and normalized to the mRNA expression level of endogenous references (*PPIA*, *RPL13* and *GusB*).

### Generation of rhinovirus 16 (RV16)-GFP

The cDNA pA16-GFP plasmid encoding RV16-GFP, which contains the rhinovirus sequence downstream of the T7 promoter, was a kind gift from Yury A Bochkov [23]. The reverse genetics technique developed by Lee et al. [24] was used to produce functional RV16-GFP virions. Briefly, for initial production, the pA16-GFP plasmid was linearized by incubation with the restriction enzyme EcI136II (EcoICRI, Thermo Scientific) at 10 U/μ purification using the QIAprep Spin MINI Prep Kit (Qiagen), the DNA was transcribed *in vitro* into RNA by T7 polymerase and then transfected into HeLa cells using Lipofectamine (Invitrogen). Twenty-four hours after transfection, the virus produced was purified and titrated by digital PCR after extraction and retrotranscription (Transcriptomic Platform, University of Bordeaux).

### Infection with RV16-GFP

At Days 4 to 5, an approximately 5 cm long bronchioid was transferred into a 32 mm diameter DMEM-containing Petri dish. The bronchioid was cannulated by a 30G needle (MicrolanceTM, Becton Dickinson, Franklin Lakes, USA) with a diameter of 300 μm, which is the order of magnitude of the inner diameter of the bronchioid. The needle was then connected to a 200 μL syringe containing DMEM (Fisher Scientific, Illkirch, France) with RV16-GFP at a concentration of 10 000 particles/μL and supplemented with 0.1% eosin (Clinisciences, Nanterre, France). The solution was manually injected into the bronchioid lumen, with visual control thanks to the eosin stain. The cultures were not washed after infection, but the opposite side of the bronchioid was “open”, so viral particles diffused out from the lumen into the medium. At 48 h post infection, the supernatants were collected and frozen at -80 °C for cytokine concentration measurements. The bronchioids were dissociated, and the cells were manually counted before being processed for FACS analysis. The volume of injection corresponds to the volume of the lumen, which was estimated to be between 1 µL and 3 µL, for 5 cm long bronchioids containing between 12 000 and 40 000 cells, with measured lumen diameters of approximately 200 µm. This allows us to calculate a multiplicity of infection (MOI) between 0.5 and 1.5.

### Cytokine production

The IFN-β, IFN-λ 1/3 and CXCL8 concentrations in the bronchioid medium were assayed 48 h post-infection by ELISA following the manufacturer’s recommendations (BioTechne). Values below the detection limit were considered zero.

### Flow cytometry

Epithelial cells in ALI culture were washed in PBS containing 0.5% bovine serum albumin (BSA, Sigma-Aldrich) and 2 mM EDTA (Invitrogen, Fisher Scientific, Illkirch, France) for 4 min at 37 °C and harvested by incubation with 0.05% trypsin-200 mg/L EDTA solution (Sigma-Aldrich) for 5 min, followed by the addition of trypsin neutralizing solution to neutralize trypsin (Lonza, Bâle, Suisse), as described previously [25]. For bronchioids, alginate dissolution was achieved by incubating the epithelial cell-laden tubes in DPBS for 20 min under gentle agitation. The cells were then mechanically and enzymatically dissociated by pipetting and by incubating the suspension with 0.05% trypsin-200 mg/L EDTA (Sigma-Aldrich) for 5 min. After centrifugation, the cell suspension was filtered through a 70-μm cell strainer (Corning) and immediately evaluated by flow cytometry to test for rhinovirus infection. Alternatively, annexin V-propidium iodide (Fisher Scientific) or DAPI (5 µg/ml, Sigma-Aldrich) was used for the detection of apoptotic and dead cells, respectively, according to the manufacturer’s instructions. For differentiation, cells were stained with anti-EpCAM-peridinin chlorophyll protein complex (PerCP)-Cy5.5 or its isotype control (see Table E2), fixed, and permeabilized using the Foxp3 buffer set (Fisher Scientific) or the IntraPrep Permeabilization Reagent Kit (Beckman Coulter). The cell pellet was then stained with anti-pan cytokeratin-FITC, anti-cytokeratin 5-Alexa Fluor 488, anti-SCGB1A1-FITC, anti-acetylated-tubulin-PE, anti-MUC5AC-Alexa Fluor 647 or their respective isotype controls (see Table E2) for 1 h. FACS data were acquired using a Canto II 4-Blue 2-Violet 2-Red laser configuration (BD Biosciences). Flow cytometry analysis was performed using Diva 8 (BD Biosciences).

### Immunostaining

The bronchioids were fixed with 4% paraformaldehyde (Sigma-Aldrich) diluted in DPBS supplemented with calcium (Sigma-Aldrich) for 30 min at room temperature and immunostained as described previously [26]. Briefly, after fixation, permeabilization and blocking steps were successively performed by incubating the bronchioids with cold DPBS supplemented with calcium containing 0.1% (vol/vol) Tween (P1379, Sigma-Aldrich) and with cold organoid washing buffer (OWB) composed of PBS containing 2 g/L BSA and 0.1% (vol/vol) Triton X-100 (Biosolve, Dieuze, France) at 4 °C for 15 min. The bronchioids were then incubated with primary antibodies diluted in OWB at 4 °C overnight with mild rocking (see Table E2). The bronchioids were washed three times with OWB for two hours each and then stained with secondary antibodies, DAPI and phalloidin diluted in OWB at 4 °C overnight, with mild rocking (see Table E2), before three supplementary washes with OWB for two hours each. Next, the bronchioids were transferred to a µ-slide 4-well chamber slide (Ibidi, Gräfelfing, Germany) and resuspended in a fructose–glycerol clearing solution composed of 60% (vol/vol) glycerol and 2.5 M fructose in dH2O or in DPBS supplemented with calcium.

### Imaging

Fixed bronchioids were imaged using a confocal Leica SP8 WLL2 microscope on an inverted DMI6000 stand (Leica Microsystems, Mannheim, Germany) using a 20X multi-immersion (NA 0.75 HC Plan Apo CS2) objective. White light laser 2 (WLL2) was used for the excitation of Alexa Fluor 647, 568 and 488 in the ranges of 650 to 750 nm, 590 to 640 nm and 500 to 550 nm, respectively. The confocal microscope was also equipped with a diode laser at 405 nm. A range of 410-480 nm was used to excite DAPI. The scanning was performed using a conventional scanner (400 Hz). The image size was 1024x1024 pixels, with a pixel size of 0.568 µm and a voxel depth of 1.040 µm. Detection was achieved with 2 internal photomultiplier tubes (PMTs) and 2 internal hybrid detectors. Fluorescence images were acquired with LAS X software (Leica). Using Fiji software, the background value was subtracted from the original image, and the image contrast in the background-corrected image was adjusted. The images were then processed with Imaris software (Oxford Instruments, version 9) for segmentation and 3D visualization. The “clipping plane” function was used to truncate the image along a given plane in 3D space.

Live imaging to analyse ciliary beating frequency was performed on an inverted Leica DMi8 microscope (Leica Microsystems, Wetzlar, Germany) equipped with a Flash 4.0 sCMOS camera (Hamamatsu, Hamamatsu, Japan). The illumination system used was a Cool LED PE-4000 (CoolLED, Andover, USA). The objective used was a HCX PL Fluotar L 40X dry 0.6 NA PH2. A 37 °C/5% CO2 atmosphere was created with an incubator box and a gaz heating system (Pecon GmbH, Erbach, Germany). Brightfield images were acquired with MetaMorph software (Molecular Devices, Sunnyvale, USA) for 2 seconds at a frame rate of 1000 images/second in 9-10 different fields of the bronchioid region. The image size was 512 x 32 pixels, with a pixel size of 0.1625 µm. The number of images used for the analysis was 1024. This results in a temporal resolution of 0.97 Hz (1000 images/second divided by 1024 images). If needed, recursive alignment of the stack of images was performed using the plugin “StackReg” of the Fiji software. Regions of interest were selected on the image generated by a maximum intensity projection using the plugin “Stardist 2D” of Fiji software (Figure E6a). The mean grey intensity was measured in each of these regions over time, and fast Fourier Transformation analysis was applied using Microsoft Excel to retrieve the different frequencies. These frequencies exhibit a discrete pattern, with an interval of 0.97 Hz between each of the possible values (Figure E6b). The main frequency was taken as the ciliary beat frequency (CBF, Figure E6b). Next, in each field, the median CBF was determined from the CBF values of the regions of interest (Figure E6c). Nine to ten fields were evaluated per donor.

### Perfusion

A 3- to 4-cm-long bronchioid was cannulated with a 30G needle (MicrolanceTM, Becton Dickinson, Franklin Lakes, USA) with a diameter of 300 μm. The needle was then connected to a microfluidic system (Fluigent, Le Kremlin-Bicêtre, France) composed of a microfluidic low-pressure generator (FLPG Plus), a microfluidic vacuum pressure-based controller (LineUp™ Push-PullTM), a Non-Invasive Flow Sensor (NIFS) for liquid or air perfusion and a reservoir with a pressurization cap (P-CAP). Air was injected into the lumen at one opened end of the bronchioid at a continuous flow rate of 0.2 mL/min. Air was released in the medium from the opposite tube end, generating visible air bubbles. Imaging was performed using an Olympus CKX53 microscope with an Olympus DP23 camera with objectives of 10X or 20X. Perfusion by air was continued for 4 hours. After the infusion was stopped, calcein (Invitrogen) was added to the culture medium at a final concentration of 1 μM, and the cells were incubated for 15 min at 37 °C.

### scRNA-seq data acquisition and analysis

A single-cell suspension was obtained from bronchioids at day 21 as described for flow cytometry analysis. Cell number and viability were determined by a Trypan blue exclusion assay before proceeding with the 10X Genomics (Pleasanton, CA) protocol for 3’ transcript capture and single-cell library preparation. Approximately 18000 cells were loaded on a 10X Chromium Next GEM Chip G. GEM generation, RT, cleanup, cDNA amplification, fragmentation, end repair & A-tail prep, and sample index tagging were performed using the Chromium Next GEM Single Cell 3’ GEM, Library & Gel Bead Kit v3.1 per the manufacturer’s instructions.

The quantification of the libraries was performed using a QubitTM 2.0 Fluorometer (Thermo Fisher), and the quality of the libraries was checked with a 2100 Bioanalyzer with a high-sensitivity DNA kit (Agilent). Libraries were sequenced using a P3 flow cell on a NextSeq2000 in 100 bp paired-end mode (Illumina) to read depths of 111,543 mean reads/cell (Patient 1) and 103,719 mean reads/cell (Patient 2).

The raw sequencing data were aligned to the GRCh38 human genome assembly (Ensembl98), and the transcripts were quantified using the CellRanger software suite (v7.1.0). Subsequently, the count matrices were loaded into R (v4.2.3), and downstream analyses were conducted using the Seurat R package (v5.0.1, [27]). Cells were filtered based on mitochondrial gene counts (<10%) and the number of genes detected per cell (>2000). Gene counts were log-normalized (NormalizeData, default parameters), and highly variable features (HVGs) were selected (FindVariableFeatures, default parameters). Batch correction was applied utilizing the Seurat reciprocal principal component analysis (RPCA) workflow with “Patient” as a covariate. A nearest neighbour graph was constructed (FindNeighbors, dims = 1:10), and a two-dimensional embedding was computed with the uniform manifold approximation and projection (UMAP) algorithm (RunUMAP, dims = 1:10). Unsupervised clustering was performed using the Louvain algorithm (FindClusters, resolution = 0.5). Cell types were annotated based on the expression of known marker genes. Ionocytes and Hillock-like cells did not cluster separately in the unsupervised clustering due to their small cell number and were selected based on UMAP coordinates, as they exhibited clearly distinct locations on the UMAP representation. Differential gene expression analysis (DGE) was performed using the Wilcoxon rank-sum test implemented in the Seurat package (FindAllMarkers, only.pos = TRUE, min.pct = 0.25), and the DGE results can be found in Table E3. Additionally, cell types were predicted with the CellTypist [28] Python package (v2.1.0) using Human_Lung_atlas.pkl as a model, with majority voting enabled. CellTypist-predicted annotations were grouped into broader categories (e.g., ‘Basal resting’ and ‘Suprabasal’ were categorized as ‘Basal’) to facilitate comparability with the manual annotations. The adjusted Rand index (ARI) was computed to assess concordance between manual annotations and CellTypist predictions.

### Primary lung tissue reference construction

Primary lung tissue scRNA-seq data along with associated cell type annotations, clinical metadata, and technical metadata were obtained from the full Human Lung Cell Atlas (HLCA) as disclosed in March 2024 (https://cellxgene.cziscience.com/collections/6f6d381a-7701-4781-935c-db10d30de293). Cells annotated as airway epithelium were extracted from the dataset. To ensure the technical coherence of the primary tissue reference with the bronchioid data produced in the present study, only cells analysed with 10X Genomics 3’ v2 or v3 assays and derived from fresh tissue were retained for reference construction. Gene counts were log-normalized, and 3000 HVGs were selected and scaled for subsequent data integration using the Seurat RPCA workflow with “dataset” as a covariate. A nearest neighbour graph was constructed, and a two-dimensional UMAP embedding was computed based on 30 RPCA dimensions.

### Neighbourhood correlation analysis

Neighbourhood graph correlation-based similarity analysis [29] was employed to compare cell states in the bronchioid model to those in primary lung tissue from distinct anatomical locations based on the common coordinate framework (CCF) established in the HLCA [30]. Cells from the primary lung tissue reference (constructed as described above) were categorized into four CCF categories: CCF 0, corresponding to the inferior turbinate; CCF 0.36-0.5, corresponding to the trachea and the main bronchi; CCF 0.72-0.81, corresponding to lobular bronchi and distal lobular airways; and CCF 0.97, corresponding to the parenchyma [30]. For the bronchioid data as well as the primary lung tissue reference CCF categories, independent neighbourhood graphs based on transcriptional similarity were constructed with the miloR package (v1.6.0) using k = 30 nearest neighbours and 30 RPCA dimensions of the respective data source [31]. The R package scrabbitr (v0.1.0) was used to calculate the mean expression profiles within each neighbourhood and to compute the Pearson correlations of expression of the 3000 HVGs from the primary lung tissue reference, excluding mitochondrial genes, between all pairs of bronchioid and CCF category neighbourhoods. The maximum correlation between each bronchioid neighbourhood and a neighbourhood in the CCF-stratified primary tissue reference was considered a measure of similarity.

## Data and code availability

The single-cell RNA-sequencing data generated in the current study are available from Zenodo under 10.5281/zenodo.10834205. The R code used for scRNA-seq data analysis has been deposited on GitHub: https://github.com/saliba-lab/Bronchioid_scRNAseq

## Results

### Formation of the tubular bronchial epithelial model from patient-derived basal cells

Basal bronchial epithelial cell cultures were obtained from 29 patients (Table 1, n= 18 non-COPD patients, n=11 COPD patients). Using the Cellular Capsule Technology, we encapsulated bronchial epithelial cells in the presence of Matrigel into tubular structures made of alginate (figure 1a). Notably, cells are injected at a concentration sufficient to allow immediate coverage of the entire alginate wall as soon as the tube is generated. Of the 29 samples, only one culture from a non-COPD patient (3%) failed to generate bronchioids due to a technical problem (figure E1). Nine bronchioids (n=4 non-COPD patients, n=5 COPD patients) were fully used before 9 days for early characterization (figure E1). Among the 19 bronchioids kept at least 11 days in culture, 6 (32%) were from patients with COPD, which is a proportion close to the composition of the enrolled patient population. Taken together, these findings show that bronchioids are generated with a high success rate and that they can be cultured for several weeks in a similar way for those derived from COPD and non-COPD patients.

Within the first 24 h, epithelial cells formed a monolayer that covered the entire inner part of the tube (figure 1b). However, as previously shown with epithelial cell lines [20], the bronchioid collapsed 1-2 days after encapsulation (figure 1b). Inhibition of actomyosin contractibility using the Rho-associated protein kinase (ROCK) inhibitor Y-27632 (10 µM) prevented epithelial detachment (figure 1b), while decreasing the level of double-phospho-myosin light chain (Thr18, Ser19; PP-MLC) (figure 1c), suggesting that cell contractility was the main driving force for shrinkage.

The dissolution of the alginate gel was successfully achieved by calcium and magnesium cation depletion for 30 minutes, allowing the generation of a cell suspension. Analysis of this suspension by flow cytometry demonstrated that the tubular structures remained stable and viable for at least 9 days (figure 1d). Thus, in subsequent experiments, bronchioids were continuously treated with Y-27632 to avoid collapse. We confirmed the epithelial nature of the cells by flow cytometry and showed that at day 1 the cells were almost all positive for EpCAM and cytokeratin, similar to differentiated bronchial epithelial cells at the ALI (Table E1, figure 2a). The mRNA levels of *CDH1* (encoding E-cadherin) and *EPCAM,* which were measured by RT-qPCR, remained stable over time (figure 2b). Confocal imaging and 3D reconstruction demonstrated the organization of bronchial cells into a tubular structure with a typical epithelial morphology and a visible lumen, reminiscent of the structure of distal bronchi (figure 2c). The presence of ZO-1 and actin fibers at cell cell contacts confirmed the presence of tight junctions (figure 2d).

**Figure 2.**
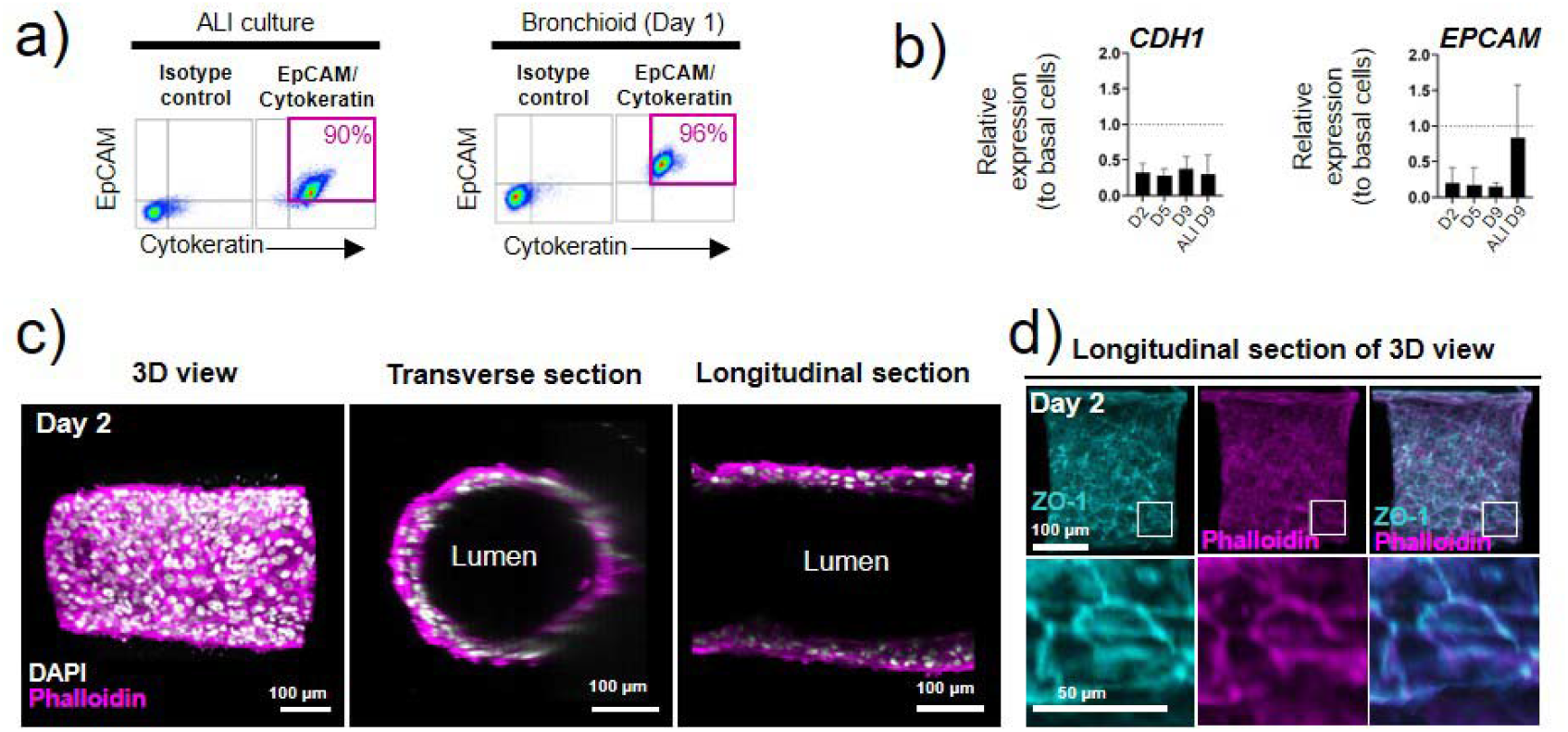
Characterization of the epithelial nature of cells in the bronchioid model. a) Dot plots representing EpCAM-PerCP-Cy5.5 fluorescence (y-axis) versus pan cytokeratin-FITC fluorescence (x-axis) of cells dissociated from 2D air-liquid interface (ALI) culture and bronchioids at day 1. The gating strategy is shown in the left panels. The percentages of EpCAM+ cytokeratin+ cells are shown in pink. b) Expression of the genes *CDH1* and *EPCAM* in bronchioids over time and in cells dissociated from 2D culture at day 9 after ALI introduction. The bronchioid and ALI culture samples were obtained from 6 and 3 different donors, respectively (mean ± standard deviation, n=6 and n=3). Gene expression was normalized to that of the housekeeping genes *PPIA*, *RPL13* and *GusB* and expressed relative to that of basal epithelial cells in 2D submerged culture. c-d) Longitudinal and transverse sections and 3D views of 3D reconstructions obtained from Z-stack confocal images of a 2-day-old bronchioid stained for F-actin (phalloidin, magenta) and nuclei (DAPI, white) (c) and F-actin (magenta), ZO-1 (cyan) and nuclei (white) (d).

### Cell differentiation and organization in the bronchioid model

RT-qPCR also revealed that the expression of the airway basal cell gene keratin 5 (*KRT5)* remained stable, and that of *TP63* progressively declined over time, whereas the expression of the club cell secretory marker secretoglobin family 1A member 1 (*SCGB1A1*) was highly upregulated (figure 3a). The expression of the ciliated cell markers *FOXJ1* and *DNAH5*, the transcription factor anterior gradient 2 (*AGR2*) and the goblet cell marker mucin 5AC (*MUC5AC*) progressively increased in culture (figure 3a). At day 9, the expression of these genes was comparable with bronchial epithelial cells at the ALI for the same time, with the exception of *EPCAM, SCGB1A1,* and *MUC5AC,* whose expression was greater, and *AGR2,* whose expression was reduced under the ALI condition (figures 2b and 3a). The cellular composition and viability were very similar in three different portions sampled along the tube (figure E2), indicating a homogenous cellular structure throughout the bronchioid. By confocal microscopy, we found that at day 11, ciliated cells were located in the internal layer lining the bronchioid lumen (figure 3b, movie E1). Most goblet cells also faced the lumen, with some MUC5AC-positive cells more externally localized (figure 3b). Positive staining for MUC5AC was also found inside the lumen, suggesting mucus secretion. Although all intermediate states of differentiation are probably not detected by simple staining, especially during the last 2 weeks in culture, flow cytometry confirmed the progressive loss of basal cells over time, with a concomitant increase in club, goblet and ciliated cells in non-COPD-derived bronchioids until day 21 (figure 3c-d). In comparison, COPD-derived bronchioids were characterized by a prominent mucous phenotype, with a proportion of goblet cells already high at day 9 (figure E3a-b). This phenotype persists until day 21, at the expense of ciliated cells whose percentage remains low. This results at day 21 in a fivefold increase and eightfold decrease in the percentage of goblet and ciliated cells, respectively, in COPD-bronchioids compared with non-COPD ones (figure E3c).

**Figure 3.**
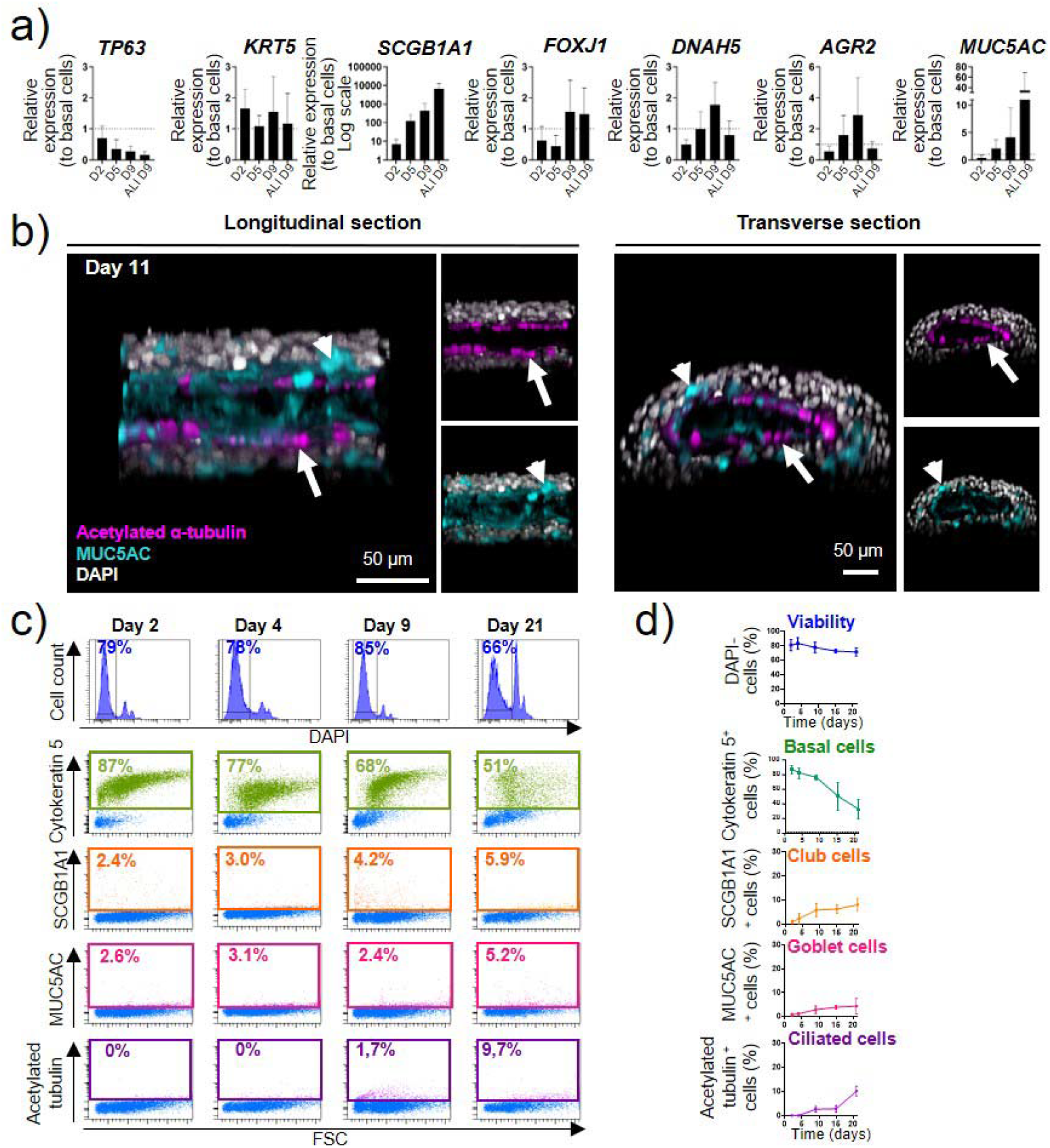
Differentiation induction of primary bronchial epithelial cells into bronchioids. a) Expression of the genes *TP63*, *KRT5*, *SCGB1A1*, *FOXJ1*, *DNAH5*, *AGR2* and *MUC5AC* in bronchioids over time and in cells dissociated from 2D culture at day 9 after ALI introduction. The bronchioid and ALI culture samples were obtained from 6 and 3 different donors, respectively (mean ± standard deviation, n=6 and n=3). Gene expression was normalized to that of the housekeeping genes *PPIA*, *RPL13* and *GusB* and expressed relative to that of basal epithelial cells in 2D submerged culture. b) Longitudinal and transverse sections and 3D views of 3D reconstructions obtained from Z-stack confocal images of 11-day-old bronchioids stained for acetylated α-tubulin (magenta), MUC5AC (cyan) and nuclei (white). Arrows: ciliated cells; arrowheads: goblet cells. c) Histograms represent representative cell counts (y-axis) versus DAPI fluorescence (x-axis) of cells dissociated from bronchioids at the indicated time points. Dot plots represent cytokeratin-5/SCGB1A1/MUC5AC/acetylated tubulin fluorescence (y-axis) versus forward side scatter (FSC, x-axis). DAPI-, cytokeratin 5+ cells, SCGB1A1+ cells, MUC5AC+ cells, and acetylated tubulin+ cells are shown in dark blue, green, orange, pink and purple, respectively. d) Percentages of DAPI-cells, cytokeratin 5+ cells, SCGB1A1+ cells, MUC5AC+ cells, and acetylated tubulin+ cells over time. The samples are from 4 different non-COPD donors (n=4).

### Single-cell characterization of non-COPD bronchioids

To further characterize the bronchioid model, single-cell RNA sequencing was conducted on bronchioids from two non-COPD patients after 21 days of culture (Methods). In total, 5,087 high-quality single cells (Patient 1 = 2,860 cells, Patient 2 = 2,227 cells) were analysed (see Methods and figure E4a) and categorized into 9 distinct populations without noticeable differences between the donors (figures 4a and E4b). These populations could be annotated based on established markers, such as basal (*KRT5*, *TP63*), secretory (*SCGB1A1*, *SCGB3A1*, *MUC5B*), goblet (*MUC5AC, CEACAM5*), deuterosomal (*FOXN4*, *CDC20B*), ciliated (*FOXJ1*, *TPPP3*), ionocyte (*FOXI1*, *CFTR*), and hillock-like (*KRT13*, *KRT14*) cells (figure 4a-b, Table E3). Additionally, populations of differentiating basal cells (*KRT5* and *AGR2*) and proliferating cells (*MKI67*) were identified. To validate the manual cell type annotations, CellTypist [28] with the Human Lung Cell Atlas (HLCA) [30] model was used for cell type prediction and demonstrated a high level of agreement (adjusted rand index = 0.691, figure E4c). Notably, all cell types, except for hillock-like cells (14 cells, 0.24%), were consistently observed in bronchioids derived from both patients, even if the proportions were not strictly identical (figures 4c and E4b). Although differentiated secretory and multiciliated lineage cells, as well as rare ionocytes and Hillock-like cells, were revealed in our bronchioid scRNA-seq data, a distinct population of *SCGB3A2*+*SFTPB*+ terminal airway-enriched secretory cells (TASCs) [32, 33] was not identified as a cluster despite the coexpression of marker genes in a few cells (figure E4d-e). Finally, to evaluate whether the bronchioid model accurately recapitulates small airways specifically at the transcriptional level, data from the HLCA and the common coordinate framework (CCF) introduced therein were utilized [30]. A primary tissue reference across four distinct anatomical regions based on CCF was assembled, comprising cells from eight datasets marked broadly as airway epithelium in the HLCA (see Methods, figure E5a-c). Bronchioid model cell states were compared to this primary tissue reference by performing a neighbourhood graph correlation-based similarity analysis ([29], see Methods). Overall, bronchioid cell states were more similar to cell states from the airways (CCF 0.32-0.5 and CCF 0.72-0.81) than to those from the nose (CCF 0) or parenchyma (CCF 0.97). Comparing the upper and lower airways, the similarity analysis results indicated that cell states from the distal airways (CCF 0.72-0.81) tended to more closely resemble bronchioid cell states (figure 4d-e). In summary, these data are consistent with the induction of distal airway fate in bronchioids, with identified cell states recapitulating distal airway epithelial cells at a global transcriptional level with high fidelity and differentiated cells localized to their corresponding *in vivo* positions.

**Figure 4.**
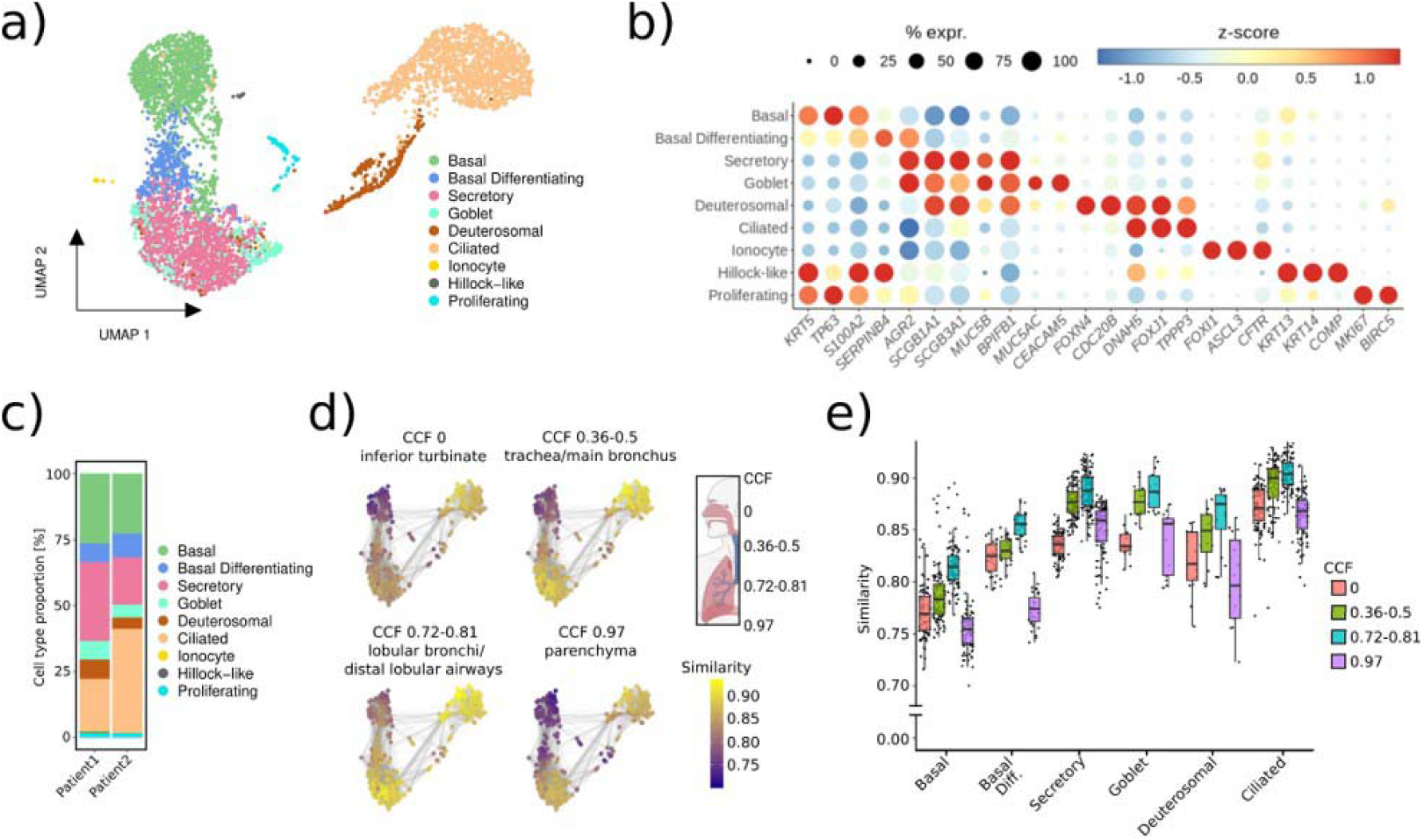
Single-cell transcriptome profiling in the bronchioid model. a) UMAP representation of scRNA-seq transcriptomic data showing the different cell types detected in 21-day-old bronchioids from 2 different non-COPD donors (*i.e.*, n=2, Patients 1 and 2). b) Dot plot showing scaled mean expression (colour) and percentage of expressing cells (dot size) of selected marker genes in the indicated cell groups. c) Relative abundance of cell types identified in bronchioids from Patients 1 and 2. d) Bronchioid neighbourhoods, positioned with respect to the UMAP embedding of each index cell, are coloured by the maximum correlation value across primary lung tissue neighbourhoods from 4 distinct anatomical locations based on the common coordinate framework (CCF) established in the HLCA [30]. e) Box plot depicting neighbourhood similarities of bronchioid neighbourhoods with primary lung tissue neighbourhoods in the respective CCF category over major cell types identified in the bronchioid scRNA-seq data. The centerline indicates the median, the box limits indicate the upper and lower quantiles, and the whiskers indicate the 1.5x interquartile range. In d–e, the bronchioid neighbourhoods were constructed from cells integrated across the 2 samples (n=2).

### Functional analysis of ciliary beating in non-COPD- and COPD-derived bronchioids

Using high-speed video microscopy [34], ciliary beating was observed in the bronchioid model (movie E2). For bronchioids generated from 4 different patients, we determined the median ciliary beat frequency (CBF) across different fields of acquisition (figure 5a-b). Overall, the median CBF values ranged from 6 Hz to 27 Hz (figure 5c), which is within the range of *in vivo* reference CBF values [35] and is also very similar to those found in bronchial epithelial strips [36], ALI cultures [37] and airway organoids [38]. Bronchioids derived from non-COPD patients were characterized by a physiologic, biphasic spectrum of beating frequencies (figure 5c), as described for control air-liquid culture [39]. In contrast, COPD-derived bronchioids exhibit a loss of the biphasic spectrum and a shift towards lower frequencies (figure 5c).

**Figure 5.**
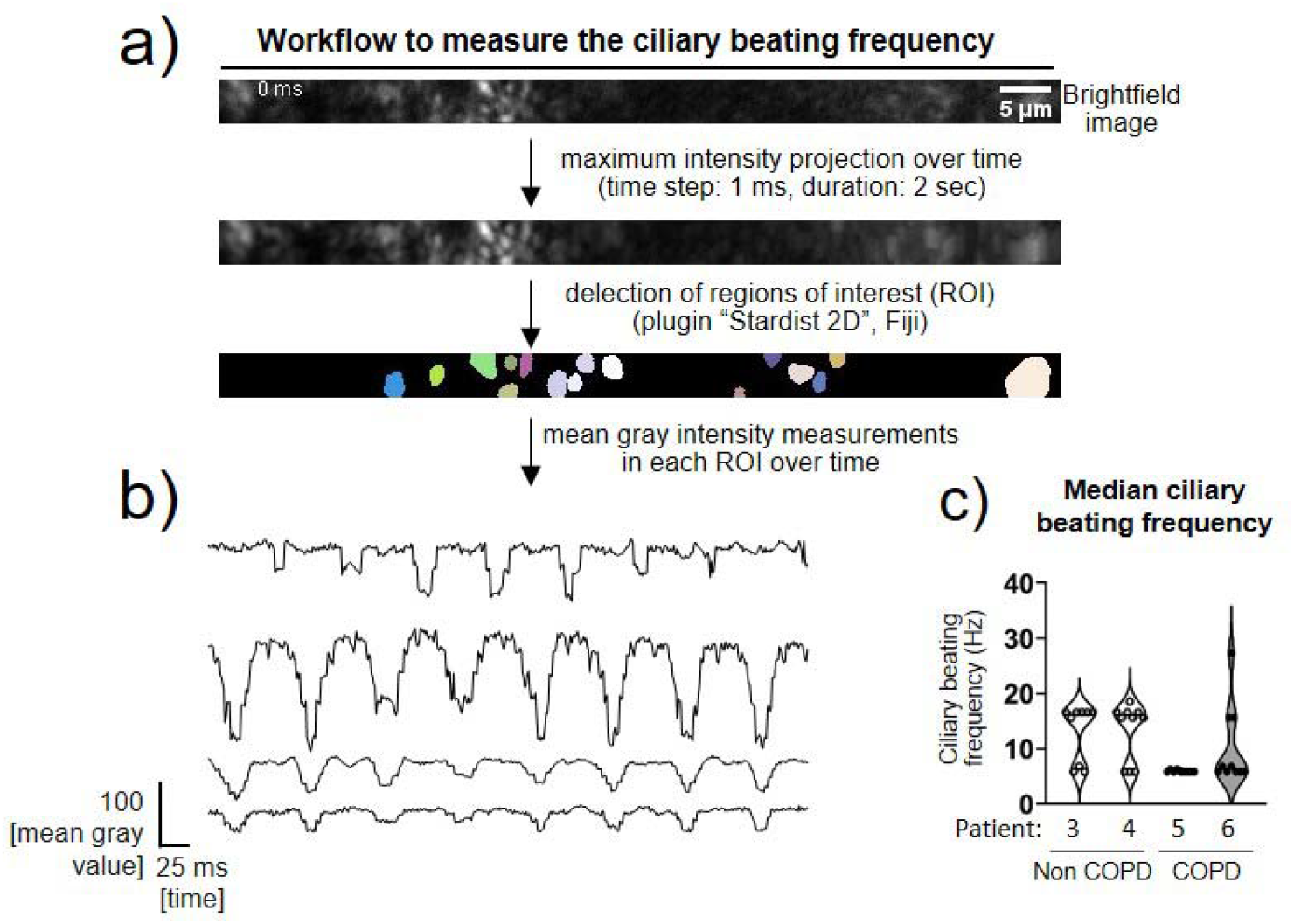
Fast Fourier transform (FFT) analysis of ciliary movement using high-speed video microscopy analysis. a) Schematic workflow for measuring ciliary beat frequency. Representative brightfield image corresponding to a part of a bronchioid at day 20. The coloured areas correspond to the regions of interest determined by the plugin “Stardist 2D” of the Fiji software on the image generated by a maximum intensity projection over time. b) Representative measurements of the mean intensity in regions of interest over time. The main frequency is determined using a fast Fourier transform in each region of interest. c) Violon plots showing the ciliary beat frequency (CBF) in Hertz (Hz) of bronchioids derived from 4 different patients (non-COPD Patients 3 and 4, COPD Patients 5 and 6). For each patient, individual values represent median CBFs, determined in 9-10 fields.

### Rhinovirus infection and epithelial response in the bronchioid model

We also used the model to test for rhinovirus infection, a major trigger of asthma and COPD exacerbation [40, 41]. To track viral infection, we injected a recombinant rhinovirus type 16 (RV16) that accommodates green fluorescent protein (GFP) expression into the lumen of the bronchioid [23] (figure 6a). Considering the calculated volume of the injection, estimated between 1 µL and 3 µL for a 5 cm long bronchioid containing between 12 000 and 40 000 cells, with measured lumen diameters of approximately 200 µm, the multiplicity of infection (MOI) was approximately 0.5 and 1.5. Forty-eight hours post-infection, 95% and 81% of the cells were GFP-positive in bronchioids generated from patients 7 and 8, respectively (figure 6b-c), suggesting infection and viral RNA replication in the vast majority of cells, in agreement with the calculated MOI. The infection was less efficient (45% of GFP-positive cells) in bronchioids generated from Patient 9 (figure 6b-c). Next, we examined the effects of RV16 on cytokine secretion, including type I (IFN-β) and type III (IFN-λ1/3) interferons and the chemokine CXCL8. RV16-GFP infection increased the levels of secreted IFN-β, IFN-λ1/3 and CXCL8, in comparison with the non-infected bronchioid condition for Patients 7 and 8 (figure 6d). No induction of interferon production or a modest increase in the CXCL8 level was observed in the bronchioids generated from Patient 9, which is in agreement with the decreased infection efficiency (figure 6d).

**Figure 6.**
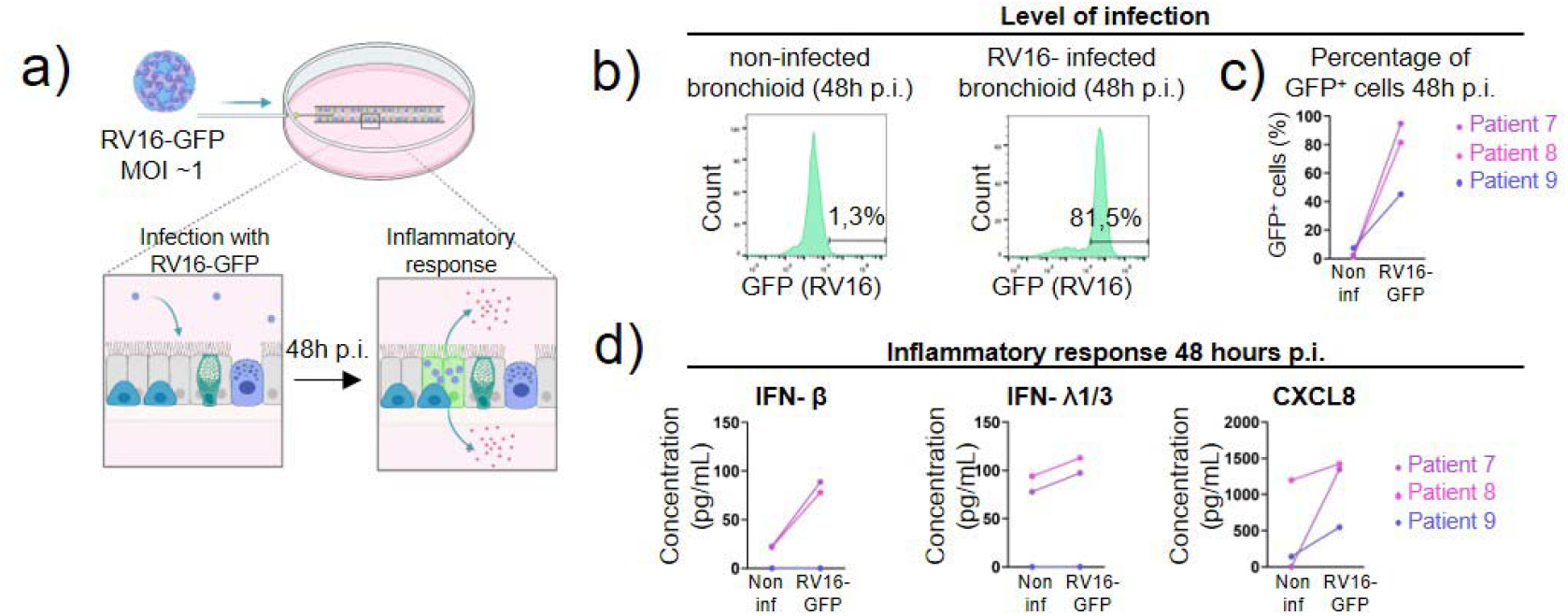
Rhinovirus infection in the bronchioid model. a) Experimental design for testing rhinovirus type 16-green fluorescent protein (RV16-GFP) infection. Multiplicity of infection (MOI). b) Histograms showing representative cell counts (y-axis) versus GFP fluorescence (x-axis) under different conditions at 48 h post-infection. The percentages indicate the number of GFP-positive cells. c) Percentages of GFP+ cells 48 h post infection, in bronchioids from 3 different donors (n=3, Patients 7, 8 and 9) under noninfected (“non inf”) and infected (“RV16-GFP”) conditions. d) IFN-β, IFN-λ 1/3 and CXCL8 concentrations in the medium from bronchioid cultures under different conditions.

### Construction of an air-conducting bronchioid model

To introduce an air-liquid interface, a 30G needle was inserted at day 3 in the lumen of the bronchioid and connected to a dedicated perfusion system (figure 7a). We injected air, as shown by a change in refringence (figure 7a, movie E3), and we maintained perfusion for 4 hours by adjusting the flow rate to match the circulating flow in the distal bronchi with an internal diameter of 400 μm, indicated that the cells *i.e.*, 0.2 mL/min. Calcein staining remained viable after 4 h of air perfusion (figure 7b). Air exposure resulted in the systematic upregulation of *SCGB1A1* in all the bronchioids perfused for 4 h (figure 7c). The expression of other genes tested showed either no variation (*TP63*, *KRT5*, *FOXJ1*, *AGR2*) or changes depending on the experiment (*DNAH5*, *MUC5AC*). Although a longer time of perfusion may be required to observe more robust effects, this indicates the potential of the ALI to affect differentiation in a bronchioid model.

**Figure 7.**
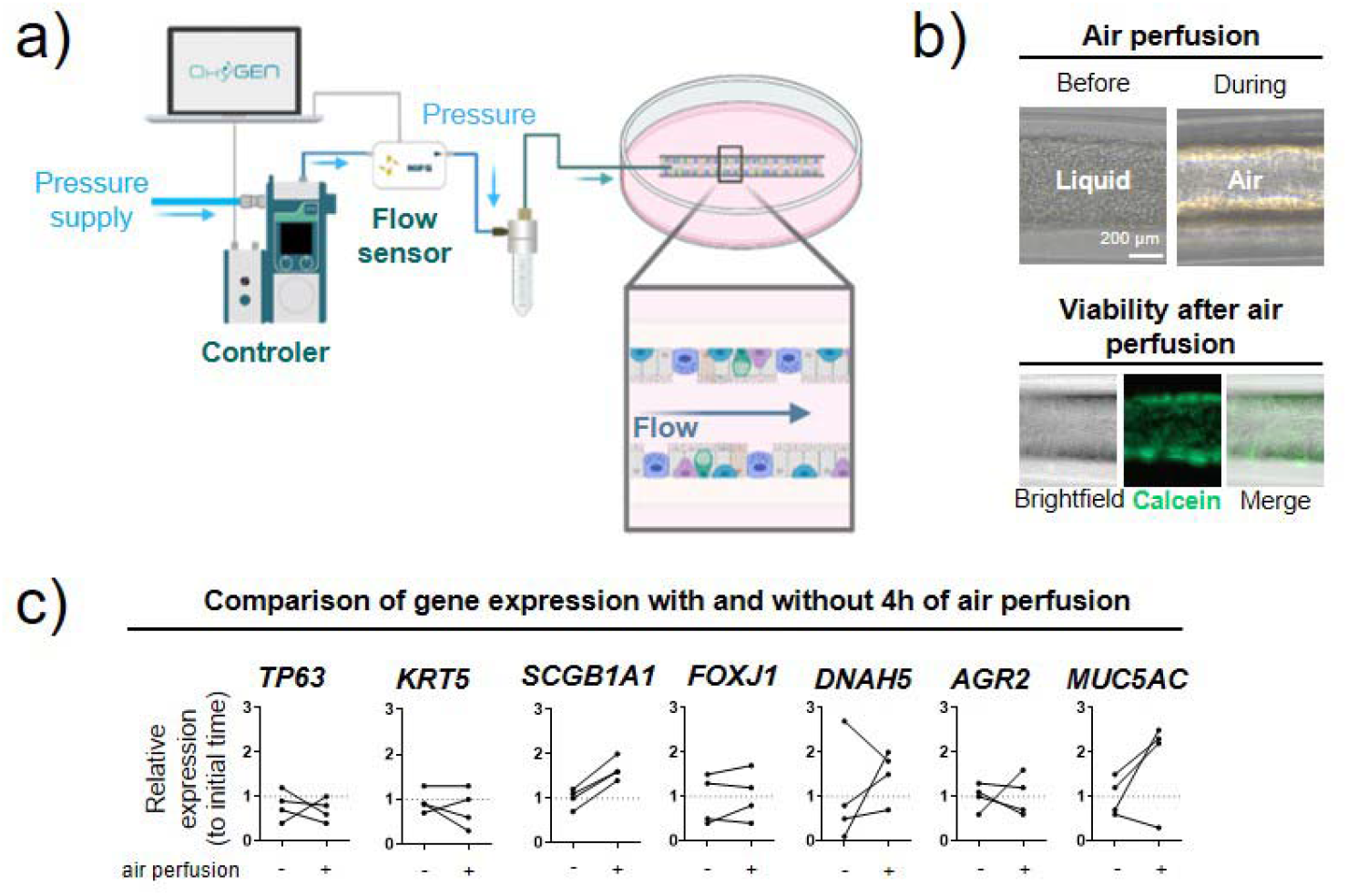
Establishment of an air-liquid interface in the bronchioid model. a) Setup for air perfusion. Created with BioRender.com. b) Upper panel: brightfield images of bronchioids before and during air perfusion. Lower panel: brightfield and fluorescence images of bronchioids after 4 h of perfusion and calcein staining. c) Comparison of gene expression (*TP63*, *KRT5*, *SCGB1A1*, *FOXJ1*, *DNAH5*, *AGR2*, *MUC5AC)* in bronchioid tissue from 4 different non-COPD donors after 4 h of perfusion (mean ± standard deviation, n=4). Gene expression was normalized to that of the housekeeping genes *PPIA*, *RPL13* and *GusB* and expressed relative to that of bronchioids before perfusion (initial time).

## Discussion

Taken together, our findings indicate that our bronchioid model offers several key advantages compared with current distal bronchial culture systems. In particular, in comparison with those of the classical 2D ALI model, the main advantages of our tubular system are the third dimension and the relevant geometry and stiffness, preventing the appearance of a nonphysiologic population by epithelial–mesenchymal transition [42] Compared with airway organoids, the bronchioid model also offers tubular geometry, lumen access for air exposure and viral infection while retaining major characteristics of the airway epithelium. This system is also compatible with multiple biological analyses, such as immunohistochemistry, flow cytometry, transcriptomics and immunoblotting. We expect this model to be instrumental in uncovering cellular mechanisms leading to airflow obstruction and for implementing efficient drug testing assays.

ROCK pathway inhibition is considered important for lung organoid generation [43] because of its antiapoptotic and proliferative effects [44]. We showed here that downregulation of MLC activity through the ROCK inhibitor Y-27632 was also required to facilitate the generation of an epithelial tubular structure that did not collapse. Importantly, the alteration of ROCK activity did not prevent cell cell adhesion or epithelial differentiation, demonstrating the key role of 3D geometry and substrate stiffness.

The incorporation of primary bronchial epithelial cells into 3D bioengineered structures is challenging, particularly due to the weak attachment of these cells to artificial substrates. The bronchioid collapse observed in the absence of ROCK inhibition may also result from insufficient adhesion to the inert alginate walls. Interestingly, tube collapse has also been observed during early lung development [45]. Thus, it is likely that there is further room for improvement in bronchioid development by favouring epithelial cell adhesion to the scaffold [7, 46] or by modulating luminal pressure [45].

By flow cytometry, we demonstrated the progressive differentiation of basal cells into club, goblet and ciliated cells. Although we cannot rule out that that differentiation may still ongoing, the bronchioid contains a subset of differentiated cells at day 21. Together with the evidence of ciliary function and mucus production, these findings indicate that the bronchioid model very likely displays fundamental mucociliary clearance properties.

At day 21, the cellular composition of the bronchioid was similar to that of airway organoids [11], with the percentage of goblet cells even slightly decreased compared to that of lung organoids. The limited set of markers used by flow cytometry did not allow us to identify all known cell types, with a substantial proportion of approximately 55% of unidentified cells at day 21. These cells likely include a majority of intermediate secretory cells, in agreement with the findings of another flow cytometry study [47]. ScRNA sequencing analysis confirmed the identification of the major bronchial epithelial cell types in proportions intermediate between those of proximal and distal airways [33], together with a high level of similarity between the bronchioid and distal airway cell types at the transcriptional level. Thus, the dimensions and geometry of the bronchioid model, which is close to a 17th generation bronchiole, as well as its general composition, make it a relevant model of the distal airway. However, the cellular composition is not strictly similar to that of distal airways, with the absence of *SCGB3A2*+ *SFTPB*+ terminal airway-enriched secretory cells (TASCs), a cell population specific to distal airways [32, 33]. This is a limitation of the current bronchioid model, which can be attributed to the proximal basal cells used in this study [32, 33] and/or to the absence of air exposure for this experiment. Indeed, our perfusion experiments indicated that the introduction of an air-liquid interface (ALI) for 4 h induced the upregulation of *SCGB1A1* expression (figure 6c). Although this requires further studies, these effects are consistent with a shift towards a distal phenotype [48] and suggest the potential of the ALI bronchioid model to fully model distal airways.

We also demonstrated that bronchioids made from cells of COPD patients might recapitulate important pathological features of COPD. First, COPD bronchioids exhibited a mucous phenotype with increased numbers of goblet cells and reduced numbers of ciliated cells. These findings are in line with those obtained in ALI cultures [39, 49] and organoid models [38] and suggest an altered differentiation trajectory in COPD bronchioids. Second, ciliary beating seems altered in COPD bronchioids, as previously shown in nasal cilia samples [50, 51] and airway organoids [38] derived from cells of COPD patients, and the loss of the biphasic distribution in ALI cultures from COPD patients [39]. Although the low number of patient replicates does not allow an accurate statistical analysis, this finding suggests the potential of using bronchioids to model respiratory diseases.

While lung organoids have been used to study infection by viruses such as syncytial virus (RSV) [11], enterovirus [52], influenza [12, 53], and acute respiratory syndrome coronavirus 2 (SARS-CoV-2) [38, 54–56], infection usually requires organoid reorientation to achieve apical-out polarity [56, 57]. Polarity reversal by suspension culture requires extracellular matrix (ECM) removal, resulting in the loss of the 3D-relevant environment and possibly affecting mechanostransduction pathways. In addition, this step takes at least 1-2 days and does not work 100% of the time. More importantly, this procedure affects cell viability in differentiated organoids, restricting its use in undifferentiated organoids. Using the bronchioid model, we demonstrated efficient rhinoviral infection through intraluminal virus injection, as well as an epithelial response. In contrast to microinjection into the centre of spherical organoids [58] or organoid mechanical disruption to expose the apical surface [59], our method does not affect the organoid wall because of the easy access to the lumen in the bronchioid model. This procedure is compatible with many types of downstream analyses, including flow cytometry and biochemical techniques. Whereas rhinovirus infection at a high dose has been previously reported to peak after 24 h in 2D ALI culture [60, 61], the high percentage of infected cells at 48 h post-infection in our study could be explained by possible reinfection, as the tube is not washed after infection. Notably, the concentration of IFN-λ 1/3 in the medium appears lower than that measured in 2D ALI culture following rhinovirus infection [60], which could be explained by the fact that the volume of medium is much greater relative to the number of cells in our settings. Overall, our results demonstrate the value of bronchioid technology for studying infectious diseases and host–pathogen interactions.

In conclusion, we generated a new workflow that allows the efficient generation of tubular structures that model distal airways and are built with epithelial cells derived from clinical samples. These bronchioids are susceptible to perfusion with liquid and, more importantly, with air to establish an ALI. As metres of tubes are generated per condition and the duration of the assay is relatively short, high-throughput methods, particularly for drug screening and testing, can be applied in the context of respiratory disease modelling.

The widespread use of these tubular models in the field of lung science requires access to dedicated technology. Notably, alginate tubes compatible with the dimensions of distal airways have been generated by different groups [18–20] and only require relatively simple co-extrusion setups, as detailed in [62]. These setups are simple in comparison to those of bioprinters or to the equipment necessary to fabricate homemade lung-on-chip systems. In addition, there are simpler methods to generate alginate tubes with larger dimensions (typically of proximal airways) [63] that are accessible with classical equipment and expertise from a biology laboratory. Thus, the adaptability and expansibility of tubular lung organoids combined with the incorporation of patient cells, together with intraluminal exposure to pathogens and particles, hold tremendous potential for the replication of specific lung pathologies. Complexification of the model with nonepithelial cells will enable us to better understand the composition and function of distal airways. Ultimately, in-depth analyses, such as single-cell transcriptomics of patient-derived bronchioids, should provide invaluable insights into disease mechanisms and facilitate screening potential therapeutics.

## Acknowledgements

We thank the study participants and the staff of the Thoracic Surgery, Respiratory, Lung Function Testing departments from the University Hospital of Bordeaux (Bordeaux, France); Isabelle Goasdoue, Isabelle Bernis, Natacha Robert, Virginie Niel, and Marine Servat from the Clinical Investigation Center for technical assistance; and Atika Zouine, Vincent Pitard, Laetitia Andrique, Vanessa Bergeron and Xavier Gauthereau for technical assistance at the VoxCell, FACSility and OneCell Facilities (CNRS UMS 3427, INSERM US 005, Univ. Bordeaux, F-33000 Bordeaux, France), Christel Poujol, Magali Mondin, Sébastien Marais and Fabrice Cordelières for help with imaging and image analysis at the Bordeaux Imaging Centre (BIC; Bordeaux, France). Microscopy was performed at the BIC, a service unit of the CNRS-INSERM, and Bordeaux University, a member of the National BioImaging Infrastructure of France supported by the French National Research Agency (ANR-10-INBS-04). AES and AML thank Deutsche Forschungsgemeinschaft DFG SFB 1583 (project number 492620490; Z2 project), and AES thanks DFG GRK2157 for funding. AES, AML and TK thank CoreUnitSysmed for providing technical support for sequencing.

## Abbreviations

ALI: Air–liquid interface
APC: Allophycocyanin
AS: Alginate solution
BEC: Bronchial epithelial cell
CCS: Core cell suspension
COPD: Chronic obstructive pulmonary disease
J3B1A: EpH4-J3B1A mammary gland epithelial cell line
EDTA: Ethylenediamine Tetra-acetic Acid
FEV1: Forced Expiratory Volume in 1 Second
FITC: Fluorescein isothiocyanate
FVC: Forced vital capacity
HeLa: Cervical cancer Henrietta Lacks cell line
HepG2: Hepatoma G2 cell line
IS: Intermediate D-sorbitol solution
MDCK: Madin-Darby Canine Kidney
MLC: Myosin light chain
MIN6m9: Mouse pancreatic beta cell line
MOI: Multiplicity of infection
OWB: Organoid washing buffer
PerCP: Phycoerythrin Peridinin Chlorophyll Protein Complex
PI: Propidium iodide
ROCK: Rho-associated protein kinase
RV16: Rhinovirus 16

## Author contributions

- conception and design (PN, GR, ID)
- data acquisition (EM, KR, VB, LA, AML, TK, AES, GR)
- data analysis (EM, KR, AML, AES, ID)
- data interpretation (EM, KR, AML, PH, MZ, RP, TT, LA, PB, PN, AES, GR, ID)
- resources (PH, TT, MT, MZ, PB)
- drafting the manuscript (EM, KR, ID)
- revision and approval of the final version (All)

## Supplemental material

The supplemental material includes Tables E1, E2 and E3; figures E1 to E6; and Movies E1, E2 and E3.

## References

1 Ghorani V, Boskabady MH, Khazdair MR, et al. Experimental animal models for COPD: a methodological review. Tob Induc Dis 2017; 15: 25.

2 Nizamoglu M, Joglekar MM, Almeida CR, et al. Innovative three-dimensional models for understanding mechanisms underlying lung diseases: powerful tools for translational research. Eur Respir Rev 2023; 32: 230042.

3 Jorissen M, Van der Schueren B, Van den Berghe H, et al. Contribution of in vitro culture methods for respiratory epithelial cells to the study of the physiology of the respiratory tract. Eur Respir J 1991; 4: 210–217.

4 Adler KB, Cheng PW, Kim KC. Characterization of guinea pig tracheal epithelial cells maintained in biphasic organotypic culture: cellular composition and biochemical analysis of released glycoconjugates. Am J Respir Cell Mol Biol 1990; 2: 145–154.

5 Dabaghi M, Carpio MB, Moran-Mirabal JM, et al. 3D (bio)printing of lungs: past, present, and future. Eur Respir J 2023; 61: 2200417.

6 De Santis MM, Bölükbas DA, Lindstedt S, et al. How to build a lung: latest advances and emerging themes in lung bioengineering. Eur Respir J 2018; 52.

7 De Santis MM, Alsafadi HN, Tas S, et al. Extracellular-Matrix-Reinforced Bioinks for 3D Bioprinting Human Tissue. Adv Mater 2021; 33: e2005476.

8 Huh D, Matthews BD, Mammoto A, et al. Reconstituting organ-level lung functions on a chip. Science 2010; 328: 1662–1668.

9 Benam KH, Villenave R, Lucchesi C, et al. Small airway-on-a-chip enables analysis of human lung inflammation and drug responses in vitro. Nat Methods 2016; 13: 151–157.

10 Benali R, Tournier JM, Chevillard M, et al. Tubule formation by human surface respiratory epithelial cells cultured in a three-dimensional collagen lattice. Am J Physiol 1993; 264: L183–192.

11 Sachs N, Papaspyropoulos A, Zomer-van Ommen DD, et al. Long-term expanding human airway organoids for disease modeling. *The EMBO Journal* John Wiley & Sons, Ltd; 2019; 38: e100300.

12 Zhou J, Li C, Sachs N, et al. Differentiated human airway organoids to assess infectivity of emerging influenza virus. Proc Natl Acad Sci U S A 2018; 115: 6822–6827.

13 Lamers MM, van der Vaart J, Knoops K, et al. An organoid-derived bronchioalveolar model for SARS-CoV-2 infection of human alveolar type II-like cells. The EMBO Journal John Wiley & Sons, Ltd; 2020; n/a: e105912.

14 Hiemstra PS, Tetley TD, Janes SM. Airway and alveolar epithelial cells in culture. Eur Respir J 2019; 54: 1900742.

15 Boecking CA, Walentek P, Zlock LT, et al. A simple method to generate human airway epithelial organoids with externally orientated apical membranes. Am J Physiol Lung Cell Mol Physiol 2022; 322: L420–L437.

16 Stroulios G, Brown T, Moreni G, et al. Apical-out airway organoids as a platform for studying viral infections and screening for antiviral drugs. Sci Rep Nature Publishing Group; 2022; 12: 7673.

17 Tan Q, Choi KM, Sicard D, et al. Human Airway Organoid Engineering as a Step toward Lung Regeneration and Disease Modeling. Biomaterials 2017; 113: 118–132.

18 Onoe H, Okitsu T, Itou A, et al. Metre-long cell-laden microfibres exhibit tissue morphologies and functions. Nature Mater Nature Publishing Group; 2013; 12: 584–590.

19 Andrique L, Recher G, Alessandri K, et al. A model of guided cell self-organization for rapid and spontaneous formation of functional vessels. Sci Adv 2019; 5: eaau6562.

20 Maechler FA, Allier C, Roux A, et al. Curvature-dependent constraints drive remodeling of epithelia. J Cell Sci 2019; 132: jcs222372.

21 Global Initiative for Chronic Obstructive Lung Disease-Internet]. Global Initiative for Chronic Obstructive Lung Disease - GOLD-cited 2023 Feb 6]. Available from: https://goldcopd.org/.

22 Trian T, Allard B, Dupin I, et al. House dust mites induce proliferation of severe asthmatic smooth muscle cells via an epithelium-dependent pathway. Am J Respir Crit Care Med 2015; 191: 538–546.

23 Olszewski D, Georgi F, Murer L, et al. High-content, arrayed compound screens with rhinovirus, influenza A virus and herpes simplex virus infections. Sci Data Nature Publishing Group; 2022; 9: 610.

24 Lee W-M, Wang W, Bochkov YA, et al. Reverse genetics system for studying human rhinovirus infections. Methods Mol Biol 2015; 1221: 149–170.

25 Maestre-Batlle D, Pena OM, Hirota JA, et al. Novel flow cytometry approach to identify bronchial epithelial cells from healthy human airways. Sci Rep Nature Publishing Group; 2017; 7: 42214.

26 Dekkers JF, Alieva M, Wellens LM, et al. High-resolution 3D imaging of fixed and cleared organoids. Nat Protoc Nature Publishing Group; 2019; 14: 1756–1771.

27 Hao Y, Stuart T, Kowalski MH, et al. Dictionary learning for integrative, multimodal and scalable single-cell analysis. Nat Biotechnol Nature Publishing Group; 2024; 42: 293–304.

28 Domínguez Conde C, Xu C, Jarvis LB, et al. Cross-tissue immune cell analysis reveals tissue-specific features in humans. Science American Association for the Advancement of Science; 2022; 376: eabl5197.

29 Ton M-LN, Keitley D, Theeuwes B, et al. An atlas of rabbit development as a model for single-cell comparative genomics. Nat Cell Biol Nature Publishing Group; 2023; 25: 1061–1072.

30 Sikkema L, Ramírez-Suástegui C, Strobl DC, et al. An integrated cell atlas of the lung in health and disease. Nat Med Nature Publishing Group; 2023; 29: 1563–1577.

31 Dann E, Henderson NC, Teichmann SA, et al. Differential abundance testing on single-cell data using k-nearest neighbor graphs. Nat Biotechnol Nature Publishing Group; 2022; 40: 245–253.

32 Kadur Lakshminarasimha Murthy P, Sontake V, Tata A, et al. Human distal lung maps and lineage hierarchies reveal a bipotent progenitor. Nature 2022; 604: 111–119.

33 Rustam S, Hu Y, Mahjour SB, et al. A Unique Cellular Organization of Human Distal Airways and Its Disarray in Chronic Obstructive Pulmonary Disease. Am J Respir Crit Care Med 2023; 207: 1171–1182.

34 Jackson CL, Bottier M. Methods for the assessment of human airway ciliary function. Eur Respir J 2022; 60: 2102300.

35 Low PM, Luk CK, Dulfano MJ, et al. Ciliary beat frequency of human respiratory tract by different sampling techniques. Am Rev Respir Dis 1984; 130: 497–498.

36 Thomas B, Rutman A, Hirst RA, et al. Ciliary dysfunction and ultrastructural abnormalities are features of severe asthma. Journal of Allergy and Clinical Immunology 2010; 126: 722–729.e2.

37 Hirst RA, Jackson CL, Coles JL, et al. Culture of Primary Ciliary Dyskinesia Epithelial Cells at Air-Liquid Interface Can Alter Ciliary Phenotype but Remains a Robust and Informative Diagnostic Aid. PLOS ONE Public Library of Science; 2014; 9: e89675.

38 Chan LLY, Anderson DE, Cheng HS, et al. The establishment of COPD organoids to study host-pathogen interaction reveals enhanced viral fitness of SARS-CoV-2 in bronchi. Nat Commun Nature Publishing Group; 2022; 13: 7635.

39 Stoleriu MG, Ansari M, Strunz M, et al. COPD basal cells are primed towards secretory to multiciliated cell imbalance driving increased resilience to environmental stressors. Thorax-Internet] BMJ Publishing Group Ltd; 2024-cited 2024 Feb 7];. Available from: https://thorax.bmj.com/content/early/2024/02/05/thorax-2022-219958.

40 Wedzicha JA. Role of Viruses in Exacerbations of Chronic Obstructive Pulmonary Disease. Proc Am Thorac Soc American Thoracic Society - PATS; 2004; 1: 115–120.

41 Message SD, Laza-Stanca V, Mallia P, et al. Rhinovirus-induced lower respiratory illness is increased in asthma and related to virus load and Th1/2 cytokine and IL-10 production. Proceedings of the National Academy of Sciences Proceedings of the National Academy of Sciences; 2008; 105: 13562–13567.

42 Greaney AM, Adams TS, Brickman Raredon MS, et al. Platform Effects on Regeneration by Pulmonary Basal Cells as Evaluated by Single-Cell RNA Sequencing. Cell Rep 2020; 30: 4250–4265.e6.

43 Kim J, Koo B-K, Knoblich JA. Human organoids: model systems for human biology and medicine. Nat Rev Mol Cell Biol 2020; 21: 571–584.

44 Watanabe K, Ueno M, Kamiya D, et al. A ROCK inhibitor permits survival of dissociated human embryonic stem cells. Nat Biotechnol Nature Publishing Group; 2007; 25: 681–686.

45 Conrad L, Runser SVM, Fernando Gómez H, et al. The biomechanical basis of biased epithelial tube elongation in lung and kidney development. Development 2021; 148: dev194209.

46 Hamilton NJI, Lee DDH, Gowers KHC, et al. Bioengineered airway epithelial grafts with mucociliary function based on collagen IV- and laminin-containing extracellular matrix scaffolds. Eur Respir J 2020; 55: 1901200.

47 Bonser LR, Koh KD, Johansson K, et al. Flow-Cytometric Analysis and Purification of Airway Epithelial-Cell Subsets. Am J Respir Cell Mol Biol American Thoracic Society - AJRCMB; 2021; 64: 308–317.

48 Yang J, Zuo W-L, Fukui T, et al. Smoking-Dependent Distal-to-Proximal Repatterning of the Adult Human Small Airway Epithelium. Am J Respir Crit Care Med 2017; 196: 340–352.

49 Gohy S, Carlier FM, Fregimilicka C, et al. Altered generation of ciliated cells in chronic obstructive pulmonary disease. Sci Rep Nature Publishing Group; 2019; 9: 17963.

50 Yaghi A, Zaman A, Cox G, et al. Ciliary beating is depressed in nasal cilia from chronic obstructive pulmonary disease subjects. Respiratory Medicine Elsevier; 2012; 106: 1139–1147.

51 Piatti G, Ambrosetti U, Santus P, et al. Effects of salmeterol on cilia and mucus in COPD and pneumonia patients. Pharmacol Res 2005; 51: 165–168.

52 van der Sanden SMG, Sachs N, Koekkoek SM, et al. Enterovirus 71 infection of human airway organoids reveals VP1-145 as a viral infectivity determinant. Emerging Microbes & Infections Taylor & Francis; 2018; 7: 1–9.

53 Hui KPY, Ching RHH, Chan SKH, et al. Tropism, replication competence, and innate immune responses of influenza virus: an analysis of human airway organoids and ex-vivo bronchus cultures. The Lancet Respiratory Medicine 2018; 6: 846–854.

54 Han Y, Duan X, Yang L, et al. Identification of SARS-CoV-2 inhibitors using lung and colonic organoids. Nature 2021; 589: 270–275.

55 Youk J, Kim T, Evans KV, et al. Three-Dimensional Human Alveolar Stem Cell Culture Models Reveal Infection Response to SARS-CoV-2. Cell Stem Cell 2020; 27: 905–919.e10.

56 Salahudeen AA, Choi SS, Rustagi A, et al. Progenitor identification and SARS-CoV-2 infection in human distal lung organoids. Nature 2020; 588: 670–675.

57 Co JY, Margalef-Català M, Monack DM, et al. Controlling the polarity of human gastrointestinal organoids to investigate epithelial biology and infectious diseases. Nat Protoc Nature Publishing Group; 2021; 16: 5171–5192.

58 Iakobachvili N, Leon-Icaza SA, Knoops K, et al. Mycobacteria–host interactions in human bronchiolar airway organoids. Molecular Microbiology 2022; 117: 682–692.

59 Lamers MM, Beumer J, van der Vaart J, et al. SARS-CoV-2 productively infects human gut enterocytes. Science 2020; 369: 50–54.

60 Veerati PC, Troy NM, Reid AT, et al. Airway Epithelial Cell Immunity Is Delayed During Rhinovirus Infection in Asthma and COPD. Front Immunol 2020; 11: 974.

61 Warner SM, Wiehler S, Michi AN, et al. Rhinovirus replication and innate immunity in highly differentiated human airway epithelial cells. Respiratory Research 2019; 20: 150.

62 Alessandri K, Sarangi BR, Gurchenkov VV, et al. Cellular capsules as a tool for multicellular spheroid production and for investigating the mechanics of tumor progression in vitro. Proc Natl Acad Sci U S A 2013; 110: 14843–14848.

63 Da Silva IA, Gvazava N, Putra Wendi I, et al. Formalin-free fixation and xylene-free tissue processing preserves cell-hydrogel interactions for histological evaluation of 3D calcium alginate tissue engineered constructs. Frontiers in Biomaterials Science-Internet] 2023-cited 2023 Aug 23]; 2. Available from: https://www.frontiersin.org/articles/10.3389/fbiom.2023.1155919.

