## Supplementary material for "A novel in vitro tubular model to recapitulate features of distal airways: The bronchioid": Online supplement

### Supplemental figures

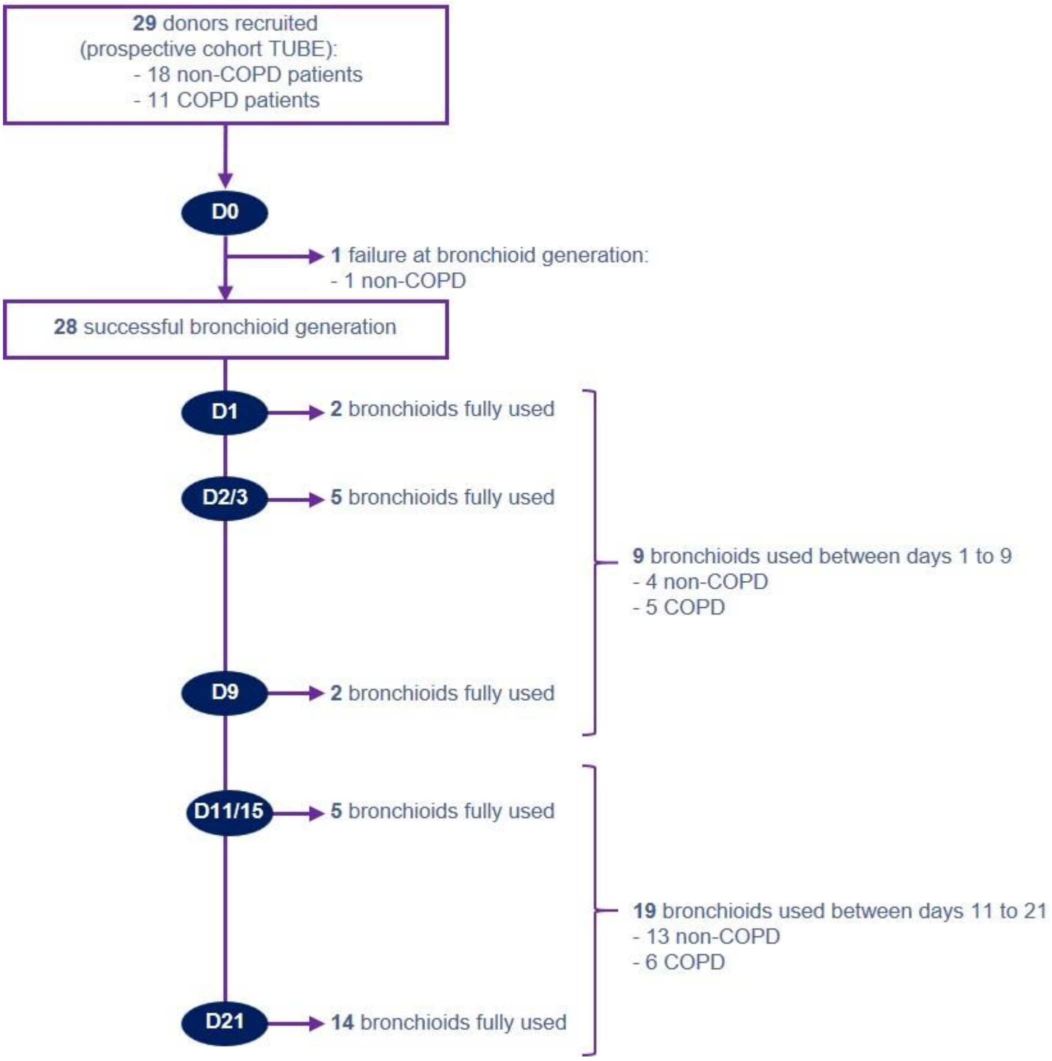


**Supplementary Fig E1. Flow diagram of sample processing for bronchioid generation and culture.** 29 patients' samples were included, 11 COPD patients with a clinical diagnosis of COPD according to the GOLD guidelines [5] and 18 non-COPD subjects with normal lung function testing (*i.e.*, $FEV_1/FVC > 0.70$). Basal bronchial epithelial cells were cultured from these samples and used to generate bronchioids, and their culture outcomes were followed. "Bronchioid fully used" means that bronchioid was kept in culture during the indicated time, and that no biological sample was left after the indicated time point.

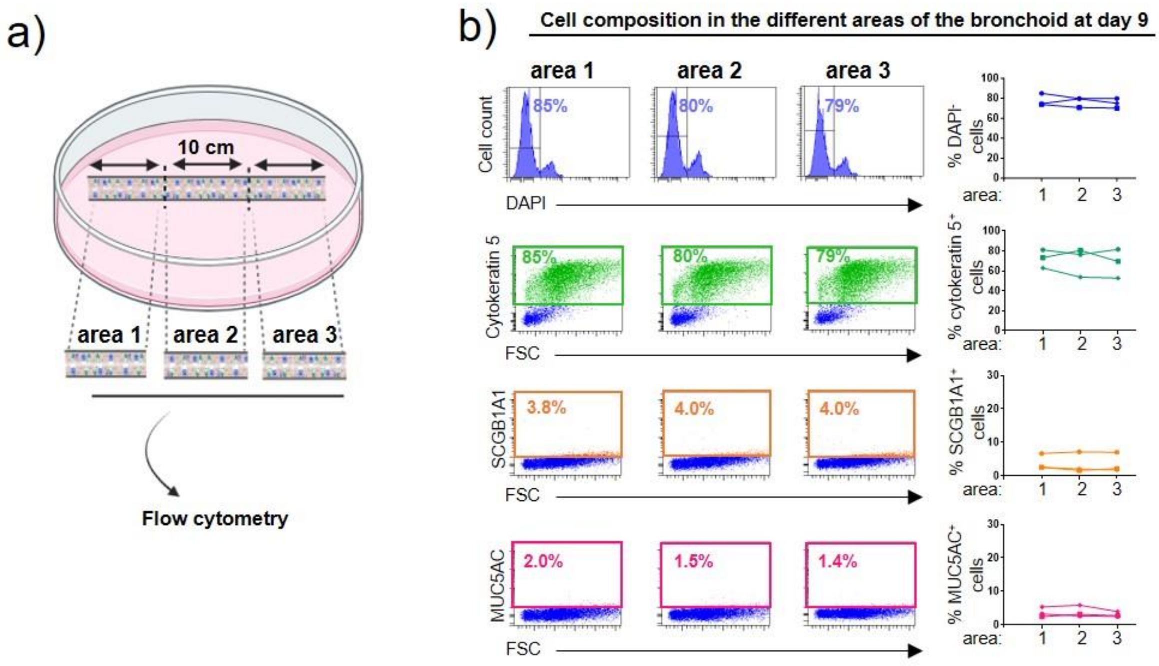


**Supplementary Fig E2. Cellular composition and viability along the tube in the bronchioid model at day 9.** a) Experimental approach to test the composition and viability in three different portions along the tube. Created with BioRender.com. b) Histograms represent representative cell count (y-axis) versus DAPI fluorescence (x-axis) of cells dissociated from bronchioids in the three portions. Dot plots represent representative cytokeratin-5/SCGB1A1/MUC5AC fluorescence (y-axis) versus forward side scatted (FSC, x-axis). DAPI$^-$, cytokeratin 5$^+$ cells, SCGB1A1$^+$ cells, MUC5AC$^+$ cells, and acetylated tubulin$^+$ cells are shown respectively in dark blue, green, orange and pink. Acetylated tubulin$^+$ cells are not represented because they are not present at day 9. c) Percentages of DAPI$^-$ cells, cytokeratin 5$^+$ cells, SCGB1A1$^+$ cells, MUC5AC$^+$ cells, and DAPI$^-$ cells over time. The samples are from 3 different donors (i.e. n=3).

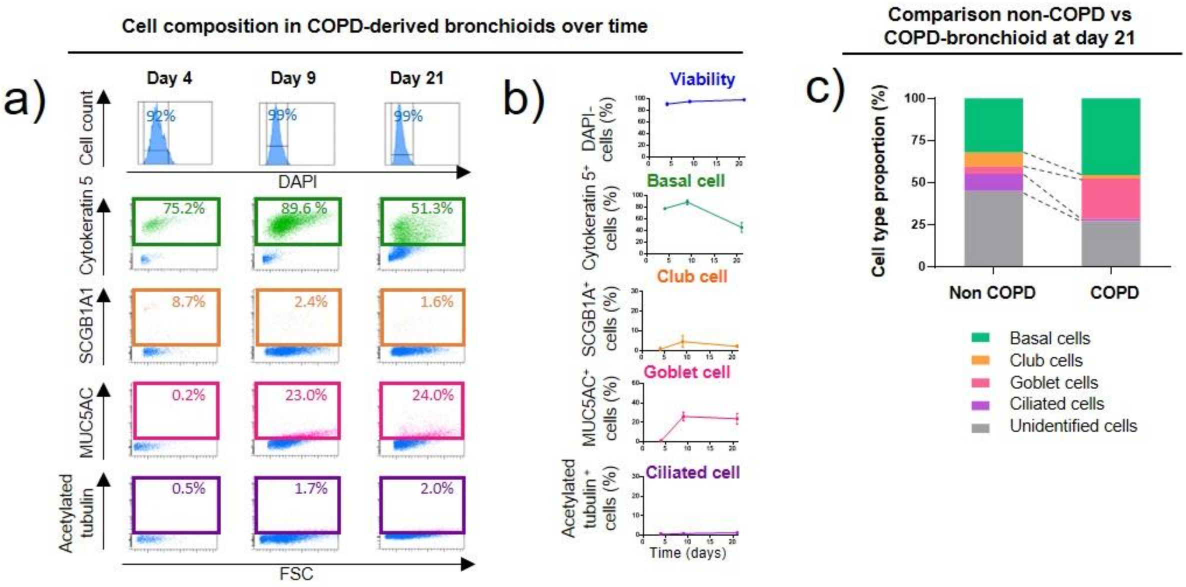


**Supplementary Fig E3. Differentiation in COPD-derived bronchioids.** a-b) The bronchioids are derived from 2 different COPD donors (i.e. n=2). a) Histograms represent representative cell count (y-axis) versus DAPI fluorescence (x-axis) of cells dissociated from bronchioids at indicated time points. Dot plots represent representative cytokeratin-5/SCGB1A1/MUC5AC/acetylated tubulin fluorescence (y-axis) versus forward side scatted (FSC, x-axis). DAPI$^{-}$cells, Cytokeratin 5$^{+}$ cells, SCGB1A1$^{+}$ cells, MUC5AC$^{+}$ cells, and acetylated tubulin$^{+}$ cells are shown respectively in dark blue, green, orange, pink and purple. b) Percentages of DAPI$^{-}$ cells, cytokeratin 5$^{+}$ cells, SCGB1A1$^{+}$ cells, MUC5AC$^{+}$ cells, and acetylated tubulin$^{+}$ cells over time. c) Comparison of relative abundance of cell types identified in 21-day-old bronchioids by flow cytometry in bronchioids from 4 different non-COPD donors (figure 3c-d) and 2 COPD donors. “Basal”, “goblet”, “club” and “ciliated” are cytokeratin-5, MUC5AC, SCGB1A1, acetylated tubulin-positive cells, respectively, “others” are unidentified cells.

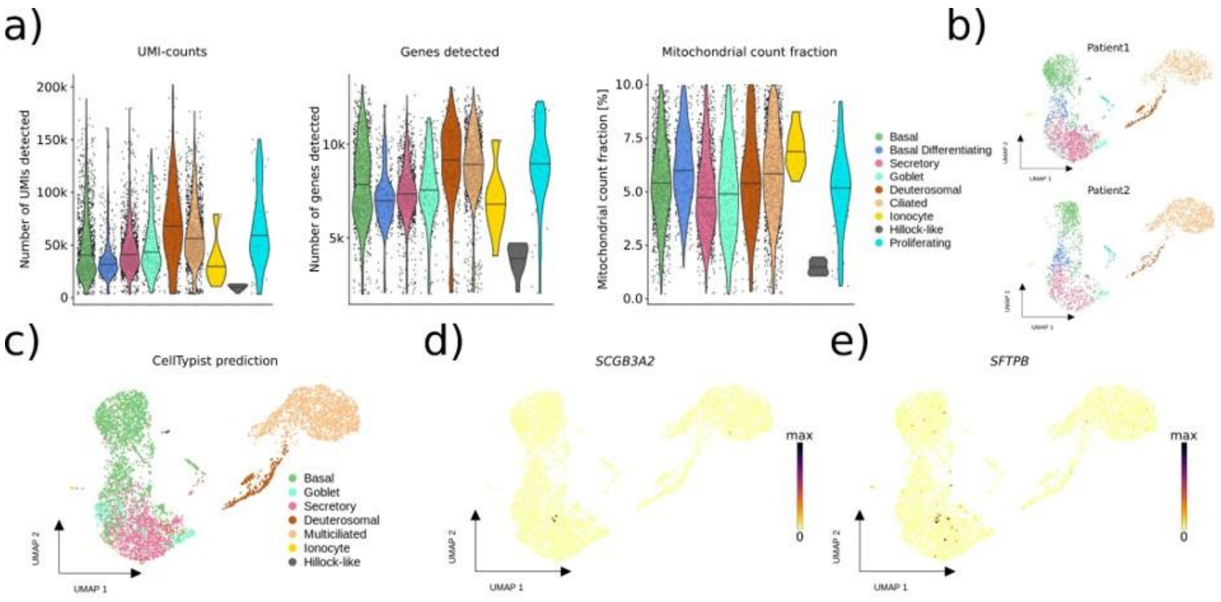


**Supplementary Fig. E4. Single cell RNA-sequencing analysis of the bronchioid model.** a) Violon plots showing the unique molecular identifier (UMI) counts (left), number of genes detected (middle) and percentage of mitochondrial count fraction (right) for each cell type detected in bronchioids from 2 non-COPD donors (i.e. n=2) after quality filtering based on UMI detected (>2000) and mitochondrial count fraction (<10%). The line in the violon plot represents the median per cell type. b) UMAP representations of scRNA-seq data as in main Figure 4a showing the different cell types detected in bronchioids generated from each of the 2 different non-COPD donors (patients 1 and 2). c) UMAP representation of the data shown in main Figure 4a colored by predicted cell types using CellTypist. d-e) UMAPs representations as in main Figure 4a showing the gene expression projection of *SCGB3A2* (d) and *SFTPB* (e).

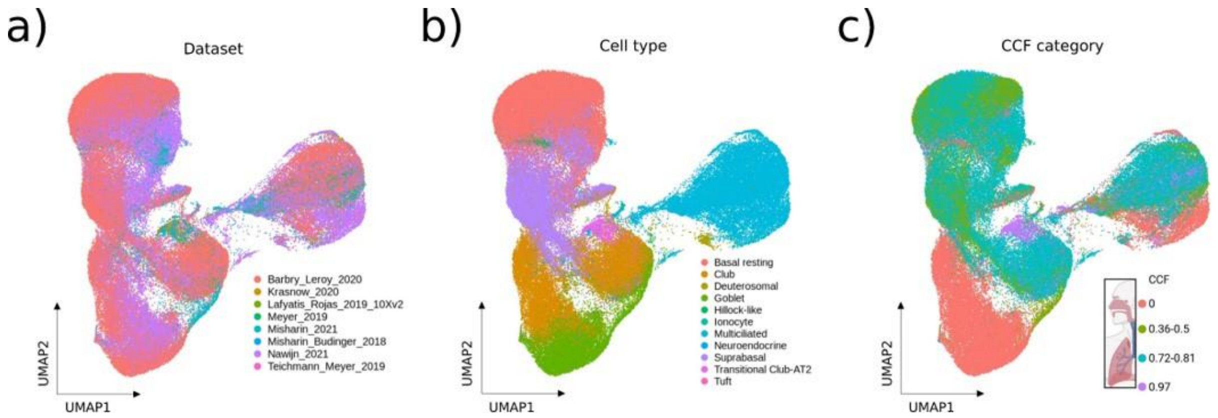


**Supplementary Fig. E5. Primary tissue reference for similarity analysis between the bronchioid model and human lung tissue.** a) UMAP representation displaying the overlap of eight scRNA-seq datasets subset to cells marked broadly as airway epithelium in the HLCA. b) UMAP representation decomposed and color-coded according to the HLCA cell type annotation in those eight aggregated datasets. c) UMAP representation showing the 4 distinct anatomical locations based on the common coordinate framework (CCF) established in the HLCA.

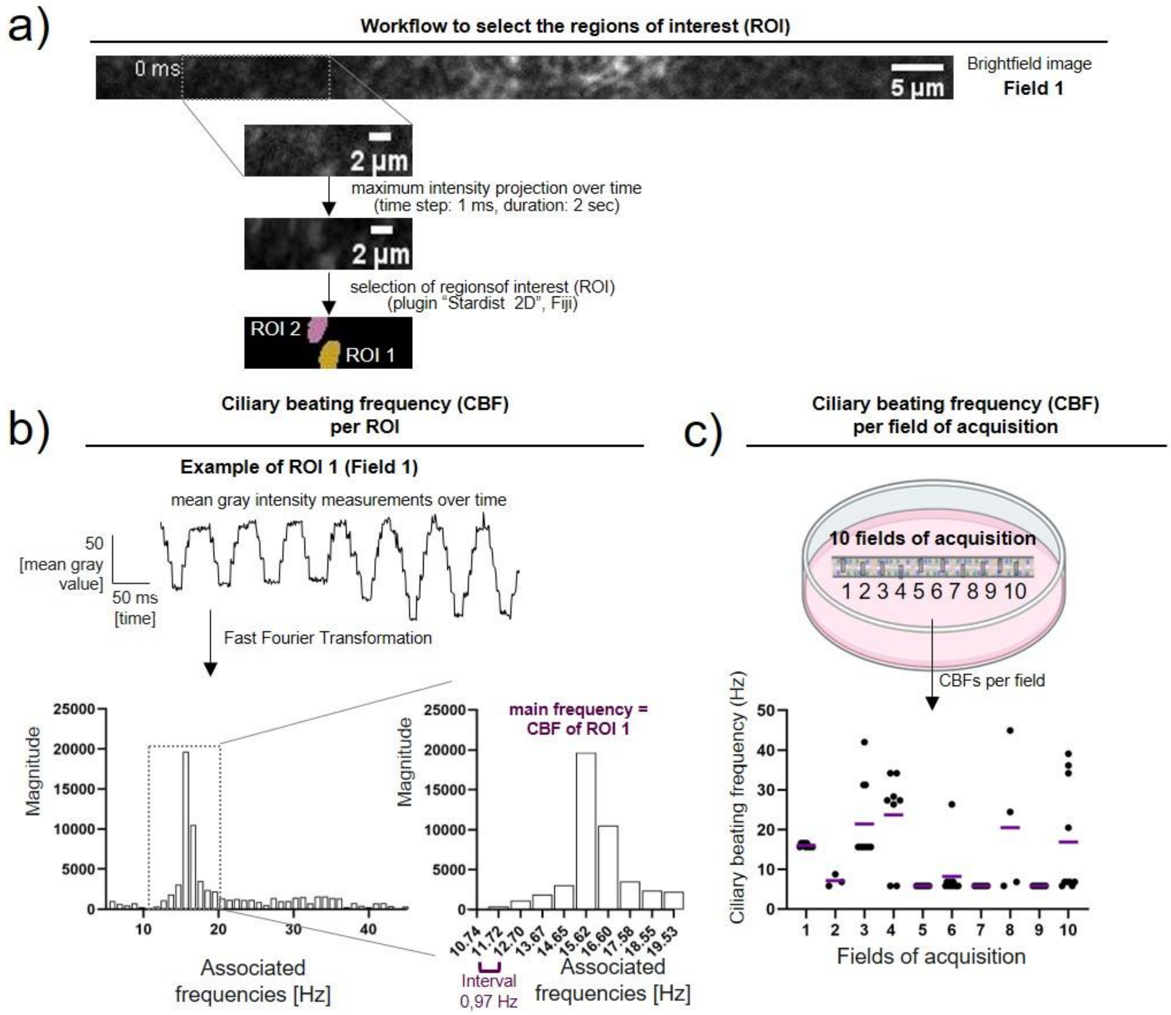


**Figure E6. Methodology for the analysis of ciliary movement using high-speed video microscopy analysis.** a) Schematic workflow to select the regions of interest (ROI) per field of acquisition, with the example of field 1. The colored areas correspond to the regions of interest determined by the plugin "Stardist 2D" of the Fiji software on the image generated by a maximum intensity projection over time. b) Measurements of the mean intensity in ROI of the field 1 over time. The frequencies are determined using fast Fourier transform and the the main frequency is taken as the ciliary beat frequency (CBF) of the ROI. c) Measurements of the CBFs in the different fields of acquisition taken on a bronchioid derived from one non-COPD patient. Individual values represent CBF in each ROI, the medians are represented as horizontal lines.

**Movie legends**

**Movie E1:** 3D reconstruction obtained from z-stack confocal images of 11-day-old bronchioid stained for acetylated α-tubulin (magenta), MUC5AC (cyan) and nucleus (white). Segmentation and 3D visualization were done using Imaris software.

**Movie E2.** High-speed video of a bronchioid at day 20. The "smart" LUT was applied to make cilia beating more visible.

**Movie E3:** Real-time video of an air-perfused bronchioid, at a flow rate of 200 µL/min. The free end of the bronchioid is imaged, the other end being connected to the perfusion system.

**Table E1: Patient characteristics for air-liquid interface (ALI) cultures**

| | Patients |
|---|---|
| n | 5 |
| COPD/non-COPD | 1/4 |
| Age (years) | 59.4 ± 20.0 |
| Sex (Men/Woman) | 2/3 |
| Current /Former/Non smokers | 0/2/3 |
| Pack years (no.) | 10.7 ± 9.0 |
| PFT | |
| FEV1 (% pred.) | 83.4± 39.8 |
| FEV1/FVC ratio (%) | 72.8± 20.4 |

Plus–minus values are means ± SD. PFT, pulmonary function test; $FEV_1$, forced expiratory volume in 1 second; FVC, forced vital capacity.

### Table E2: Resources Table – Antibodies

| Reagent type | Designation | Source or reference | Identifiers | Additional information |
|---|---|---|---|---|
| Antibody | Anti-Myosin Light Chain 2 (D18E2) (rabbit monoclonal) | Cell Signaling | Cat. #:8505 | Western Blot (1:1000) |
| Antibody | Anti-Phospho-Myosin Light Chain 2 (Thr18/Ser19) (rabbit monoclonal) | Cell Signaling | Cat. #:3674 | Western Blot (1:1000) |
| Antibody | Goat Anti-Rabbit IgG Antibody (H+L) | Vector Laboratories | Cat. #:PI-1000 | Western Blot (1:2500) |
| Antibody | Anti-EpCAM-PerCP-Cy5.5 | Sony | Cat. #:2221065 | Flow cytometry (3:100) |
| Antibody | Iso EpCam-PerCP-Cy5.5 | Sony | Cat. #:2600750 | Flow cytometry (3:100) |
| Antibody | Anti-pan cytokeratin-FITC | Invitrogen | Cat. #:MA5-28561 | Flow cytometry (3:100) |
| Antibody | Iso pan cytokeratin-FITC | Sony | Cat. #:2600690 | Flow cytometry (3:100) |
| Antibody | Anti-cytokeratin 5-Alexa 488 | Abcam | Cat. #:ab193894 | Flow cytometry (3:100) |
| Antibody | Iso cytokeratin 5-Alexa 488 | Abcam | Cat. #:ab199091 | Flow cytometry (3:100) |
| Antibody | Anti-SCGB1A1-FITC | Santa Cruz | Cat. #:sc-365992 | Flow cytometry (3:100) |
| Antibody | Iso SCGB1A1-FITC | Santa Cruz | Cat. #:sc-2855 | Flow cytometry (3:100) |
| Antibody | Anti-Mucin 5AC-Alexa 647 | Novus Bio. | Cat. #: NBP2-32732AF647 | Flow cytometry (3:100) |
| Antibody | Iso Mucin 5AC- Alexa 647 | Novus Bio. | Cat. #: IC002R | Flow cytometry (3:100) |
| Antibody | Anti-α-acetylated tubulin-PE | Cell Signaling | Cat. #:18276S | Flow cytometry (3:100) |
| Antibody | Iso α-acetylated tubulin-PE | Cell Signaling | Cat. #:5742S | Flow cytometry (3:100) |
| Antibody | Anti-ZO-1 (mouse monoclonal) | Invitrogen | Cat. #:33-9100 | IF (1:50) |
| Antibody | Anti-Mucin 5 AC (rabbit monoclonal) | Santa Cruz | Cat. #:sc-20118 | IF (1:100) |
| Antibody | Anti-α-acetylated tubulin (mouse monoclonal) | Santa Cruz | Cat. #:sc-23950 | IF (1:100) |
| Antibody | Donkey anti-Rabbit IgG Alexa Fluor™ Plus 488 | Invitrogen | Cat. #: A32790 | IF (1:100) |
| Antibody | Goat anti-Mouse IgG Alexa Fluor™ 568 | Life Technologies | Cat. #: A11004 | IF (1:100) |
| Dye | Phalloidin- iFluor 647 | Abcam | Cat. #:ab176759 | IF (1:500) |
| Dye | DAPI | Sigma-Aldrich | Cat. #:D9542 | IF (1:1000) |

**Table E3: differential expression analysis results between cell types identified in the bronchioids (excel file)**
